# Single-cell morphometrics reveals ancestral principles of notochord development

**DOI:** 10.1101/2020.07.08.193813

**Authors:** Toby G R Andrews, Wolfram Pönisch, Ewa Paluch, Benjamin J Steventon, Elia Benito-Gutierrez

## Abstract

During development, embryonic tissues are formed by the dynamic behaviours of their constituent cells, whose collective actions are tightly regulated in space and time. To understand such cell behaviours and how they have evolved, it is necessary to develop quantitative approaches to map out morphogenesis, so comparisons can be made across different tissues and organisms. With this idea in mind, here we sought to investigate ancestral principles of notochord development, by building a quantitative portrait of notochord morphogenesis in the amphioxus embryo – a basally-branching member of the chordate phylum. To this end, we developed a single-cell morphometrics pipeline to comprehensively catalogue the morphologies of thousands of notochord cells, and to project them simultaneously into a common mathematical space termed morphospace. This approach revealed complex patterns of cell-type specific shape trajectories, akin to those obtained using single-cell genomic approaches. By spatially mapping single-cell shape trajectories in whole segmented notochords, we found evidence of spatial and temporal variation in developmental dynamics. Such variations included temporal gradients of morphogenesis spread across the anterior-posterior axis, divergence of trajectories to different morphologies, and the convergence of different trajectories onto common morphologies. Through geometric modelling, we also identified an antagonistic relationship between cell shape regulation and growth that enables convergent extension to occur in two steps. First, by allowing growth to counterbalance loss of anterior-posterior cell length during cell intercalation. Secondly, by allowing growth to further increase cell length once cells have intercalated and aligned to the axial midline, thereby facilitating a second phase of tissue elongation. Finally, we show that apart from a complex coordination of individual cellular behaviours, posterior addition from proliferating progenitors is essential for full notochord elongation in amphioxus, a mechanism previously described only in vertebrates. This novel approach to quantifying morphogenesis paves the way towards comparative studies, and mechanistic explanations for the emergence of form over developmental and evolutionary time scales.

## INTRODUCTION

A major challenge in biology is to understand how individual cells coordinate their behaviours during embryogenesis to generate tissues of the correct geometry and size, and how these behaviours are modified through evolution to generate morphological novelty. The notochord is a pivotal case study in this context. It is a defining feature of the chordate body plan with diverse contributions to axial development. The notochord also has a simple geometry, as an elongate rod of mesodermal tissue occupying the embryonic axial midline (Stemple, 2005). During its development, the notochord exerts essential roles in body plan formation, including contributions to axis elongation and mechanical stabilisation of the body axis (Stemple 2005; Segade et al., 2016; Xiong et al., 2018), and the secretion of organising signals that provide dorsoventral patterning information to the adjacent neural tube and somites (Placzek et al., 1991; Pourquié et al., 1993; Yamada et al., 1993). Once formed, the notochord also provides structural support to the embryo prior to the development of a complete skeletal system. With central roles in body plan morphogenesis, innovation of the notochord likely imposed radical change in the pattern and geometry of chordate embryos. However, while notochord development has been well characterised in olfactores (ascidians + vertebrates), the cell behaviours responsible for its formation in the first chordates remain unknown. The invertebrate chordate amphioxus, representing the most basally branching chordate subphylum (Delsuc et al., 2006), provides a unique opportunity to infer which morphogenetic principles might be ancestrally linked to notochord development, and which of these represent species-specific traits.

In vertebrates and ascidians, the notochord develops from a loosely-packed field of mesodermal progenitors (termed chordamesoderm) that progressively organise into an elongated rod of tissue by changing their shapes and spatial organisation. Notochord cells actively change their shape, and crawl between adjacent neighbours to intercalate and generate a single-file row. As a result, neighbouring cells are forced apart along the anteroposterior (AP) axis, resulting in tissue extension (Glickman et al., 2003; Munro and Odell, 2002; Shih and Keller, 1992). This process is termed convergent extension, in which tissue length is established at the expense of width (Keller et al., 2000). To variable degrees in each of the species studied, convergent extension synergises with cell growth and proliferation to define total notochord length. This ranges from ascidians, where notochord morphogenesis occurs in a population of exactly 40 post-mitotic cells (Miyamoto and Crowther, 1985; Veeman and Smith, 2013), to amniotes that extensively elongate the notochord primordium after gastrulation through proliferation of posterior axial progenitors (Cambray and Wilson, 2002; Catala et al., 1996; Selleck and Stern, 1991). Intercalation is generally followed by vacuolation of individual notochord cells, which expands their volume and increases notochord length and rigidity (Adams et al., 1990; Bancroft and Bellairs, 1976; Ellis et al., 2013).

In amphioxus, the chordamesoderm is specified at the dorsal midline of the archenteron, the primitive gut cavity formed during gastrulation (Zhang et al., 1997). After gastrulation, the chordamesoderm evaginates to generate a longitudinal groove, and cells on either side interdigitate. This establishes a single file row of cells that ultimately stabilises in a trilaminar arrangement along the dorsoventral (DV) axis, described as a central row of flattened cells in a stack-of-coins pattern, flanked dorsally and ventrally by single-file rows of rounded Müller cells (Conklin, 1934). The central stack-of-coins is a feature shared with other chordates. In contrast, the Müller cells are unique to amphioxus. While their function is unknown, roles have been proposed in secretion of the notochord sheath and mechanical axial support (Bočina and Saraga-Babić, 2006; Holland and Holland, 1990; Flood, 1975). During amphioxus development, little proliferation has been reported in the notochord, except for cells at its posterior tip (Holland and Holland, 2006). Cell growth has also been described, and attributed to vacuolation (Hatschek, 1893). However, beyond these studies, a detailed understanding of the cellular behaviours collectively contributing to the formation and elongation of the amphioxus notochord remains lacking (Annona et al., 2015). With a simple morphology, small size and optical transparency, the amphioxus embryo is an ideal system for image-based quantitative analysis of individual cell shape changes and how their behaviours correlate with tissue shape change through development.

To build a complete picture of cell shape changes during amphioxus notochord development, we use large-scale morphometric techniques, that have been previously used to compare biological shapes ranging from single cells in culture to entire organisms. Approaches for shape quantification have included standard geometric measures (Mingqiang et al., 2008; Pincus and Theriot, 2007; Tassy et al., 2006), landmark-based morphometrics (Watanabe et al., 2019; Webster and Sheets, 2010), and Fourier descriptors (Boehm et al., 2011; Medyukhina et al., 2020; Tweedy et al., 2013). The resulting high-dimensional datasets are often visualised in lower-dimensional spaces, termed morphospaces, by applying dimensionality reduction approaches such as Principal Component Analysis (PCA) (Bhullar et al., 2012; Ruan et al., 2020; Tweedy et al., 2013). In morphospace, short distances between specimens reflect morphological similarity between different shapes, whereas large distances represent disparity. These patterns can expose specific transitions in form over developmental and evolutionary timescales (Morris et al., 2019; Yin et al., 2014; Young et al., 2014). Here, we reason that this approach can be used to define trajectories of cell shape change underpinning tissue morphogenesis, when applied to morphometric data for thousands of cells, from embryos at different developmental stages, across the whole spatial extent of a developing tissue. This analysis offers a systems-level view of tissue morphogenesis. From here on, we refer to such a descriptions of developmental transformations in cell shape as ‘trajectories’, akin to the description of cell state trajectories in single cell transcriptomic analyses.

Here, we define patterns of cell shape change for all cells of the amphioxus notochord during its elongation, by projecting them into a single-cell morphospace. We find that cells in morphospace cluster into a series of branching cell type-specific trajectories, which all emerge from a common progenitor morphology. Since our approach also captures each cell’s position within the embryo, we further show that cells at different positions along the AP axis progress towards common morphologies at different times, and through different morphogenetic paths. We next make use of geometric modelling, applied to mean progenitor cell shapes, to determine how specific geometric transformations contribute to the global cell shape transitions measured along each trajectory. This analysis suggests that volumetric growth enables convergent extension in the notochord in two steps: Firstly, by counterbalancing loss of AP length as cells increase their surface area during intercalation. Secondly, by directly increasing cell AP length after they have intercalated and become distributed along axial midline. Finally, we use cell labelling strategies and pharmacological cell cycle arrest to test the role of cell division in notochord elongation and expose a novel role for posterior progenitors in regulating the number of cells available for each morphogenetic trajectory. Overall, our approach reveals the notochord of amphioxus to be more morphologically complex and heterogenous than previously thought, and identifies a basic repertoire of morphogenetic processes ancestrally linked to notochord development.

## RESULTS

### A single-cell morphospace captures branching trajectories of shape differentiation specific to cell type

In amphioxus, notochord formation involves a complete restructuring of cellular organisation from a seemingly disorganised group of rounded cells at the 6 somite stage (ss), to a regular trilaminar array at the 14ss, with central cuboidal cells in a stack-of-coins pattern sandwiched between two rows of Müller cells (*Fig. 1a, insets*) (Hatschek, 1893; Conklin, 1924). Assuming that cells gradually transition towards their final morphologies, cell shapes in fixed specimens will reflect snapshots of differentiation. With this logic, we sought to reconstruct dynamic trajectories of single-cell shape change by assembling 3D single-cell morphometric data into an active developmental morphospace. To achieve this, we built a dataset of manually-segmented notochord cells at successive stages of elongation, using specimens stained with phalloidin to mark cortical actin from 6-14ss (*Fig. 1a, b*). In total 3,796 cells were segmented, from 15 notochords across 5 developmental stages. We quantified geometrical parameters of each cell that include surface area, volume, 3D long axis orientation, cross-sectional area, cell spread, smoothness, oriented anisotropy, cuboidness, sphericity, flatness, and nuclear displacement from centre of mass (*Fig. 1c;* itemised in *Materials and Methods*). To identify the major axes of shape variation between cells, we then performed a principal component analysis (PCA) on these measurements. In doing so, we found that 86.2% of cell shape variability was explained by 5 eigenvectors (*Fig. S1a*); anisotropic elongation along the AP axis (PC1, 36.7% variation; *Fig. S1c*), transverse elongation and orientation (PC2, 20.9% variation; *Fig. S1d*), volume and surface smoothness (PC3, 14.1% variation; *Fig. S1e*), surface convolution (PC4, 9.7% variation; *Fig. S1f*), and nuclear displacement from the centre of homogenous mass (PC5, 4.8% variation; *Fig. S1g*) (*Fig. 1c, S1*). These components offered a framework to visualise geometric variation in low-dimensional space.

**Figure 1.**
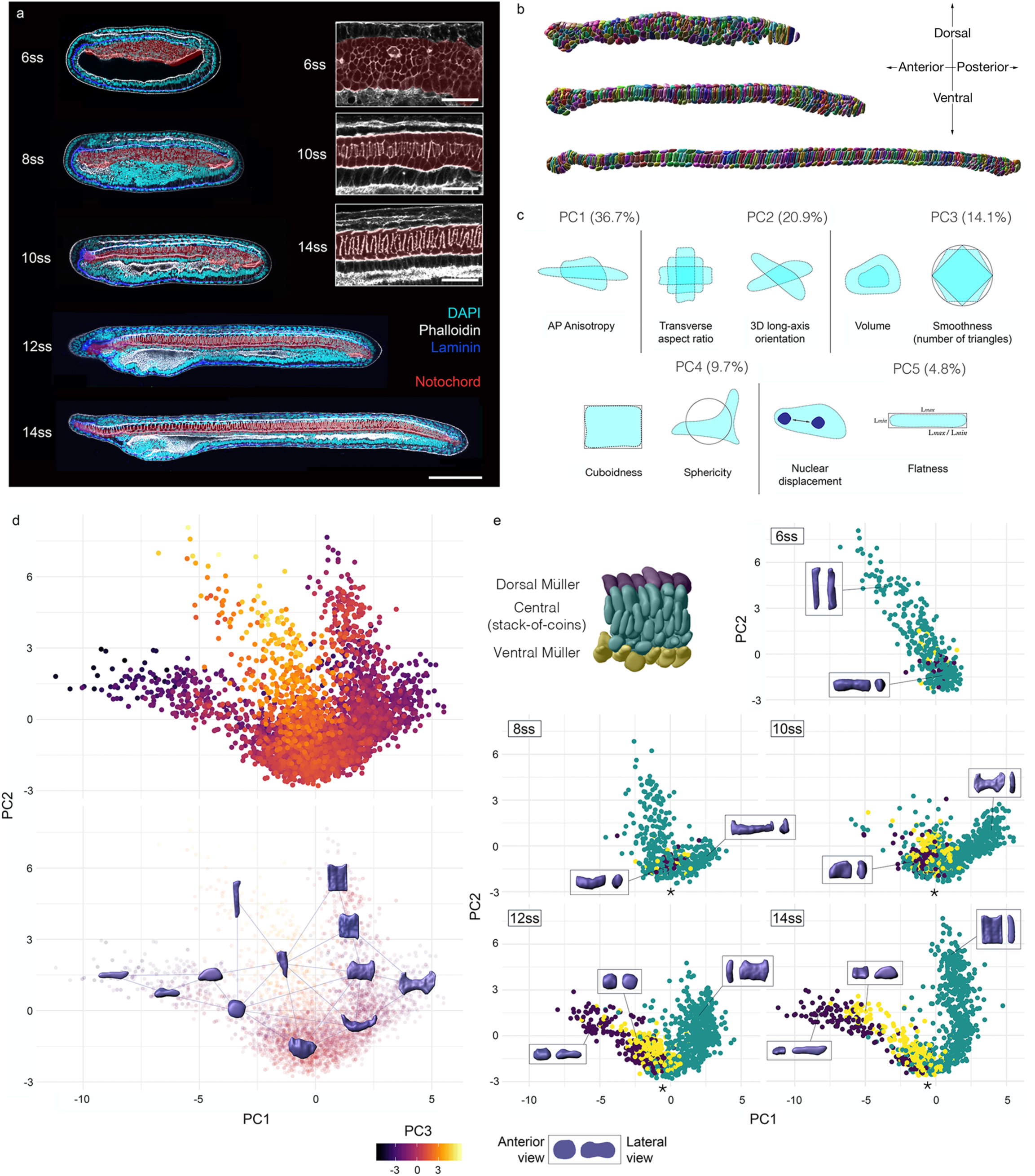
A single-cell morphospace captures notochord cell shape diversity. (a) Successive stages of amphioxus embryo spanning axis extension, stained for phalloidin, and immunostained for laminin. (b) Fully segmented notochords from the 8-, 10, and 12-somite stages. (c) Major geometric correlates of the first five principal components. (d top) Active notochord morphospace for all cells and stages, with colour code for PC3. (d bottom) Notochord morphospace illustrated with representative surfaces from segmentation data. (e) Morphospace filtered by developmental stage and colour-coded for position along the DV axis, where Müller and stack-of-coins cell types become resolved. Representative cells for each domain shown in inlays, in anterior (left) and lateral (right) views. 3,796 individually-segmented cells, 5 stages, 3 embryos per stage.

We next plotted cells against the first three PCs to construct a developmental morphospace. In morphospace, cells organised in a continuous and highly-structured branching pattern (*Fig. 1d*), with three primary branches connecting to the diversity of cell shapes that emerge during notochord development (*Fig. 1d*). This pattern was highly reproducible across the three embryos analysed per stage (*Fig. S2*). To ask how cells traverse this morphospace during their development, we subdivided the data by developmental stage. At 6ss, all cells occupied the central branch of morphospace, with tall DV-elongated morphologies (*Fig. 1e, 6ss*). Over time, these morphologies were lost, as cells flowed from the central branch into a bifurcation at the base of the plot (at the level of -1.5 on PC1) to form two distinct trajectories of shape change (*Fig. 1e, asterix*). Although cells from older specimens populated more distal positions in morphospace with respect to this bifurcation, a continuum persisted at each stage between immature and differentiated states. By additionally categorising cells based on relative DV position, we found the bifurcation event to yield independent trajectories for the two main cell types in the notochord: the central cells and Müller cells (*Fig. 1e, 10ss – 14ss*). While the Müller cells become elongated along the AP axis at the expense of their transverse area (decreasing on PC1), the central cells spread out on a transverse plane at the expense of their length (increasing on PC1). The trajectory followed by Müller cells was further divided in sub-trajectories for the dorsal and ventral rows, based on distinctive levels of AP anisotropy (*Fig. 1e, yellow and blue points*). Overall, our approach reveals a progressive diversification of cell morphology during notochord development, in which a common progenitor morphology is remodelled along diverging shape trajectories to generate a predictable diversity of cell morphologies.

### Single-cell morphometrics highlights two major shape transitions in central notochord cells

From this global view of shape differentiation, we next used our morphometric approach to define major transitions in cell shape specific to the central layer of notochord cells – its most conserved component. To infer how central cell shapes change over developmental time, we selected cells at a specific position in the notochord (the 50% level of the AP axis), and mapped their distribution in morphospace over successive developmental stages (*Fig. 2a, b*). Collectively, we found central cells from the 50% level to define a continuous trajectory of shape change through morphospace (*Fig. 2b*). Between 6ss and 10ss, cells increased on PC1 and PC2, and decreased on PC3. At 10ss, the trajectory changed direction such that cells then declined on PC1, and increased on PC2 and PC3. This trajectory structure suggested that central notochord cells experience two major morphological transitions during their differentiation (*Fig. 2b*).

**Figure 2.**
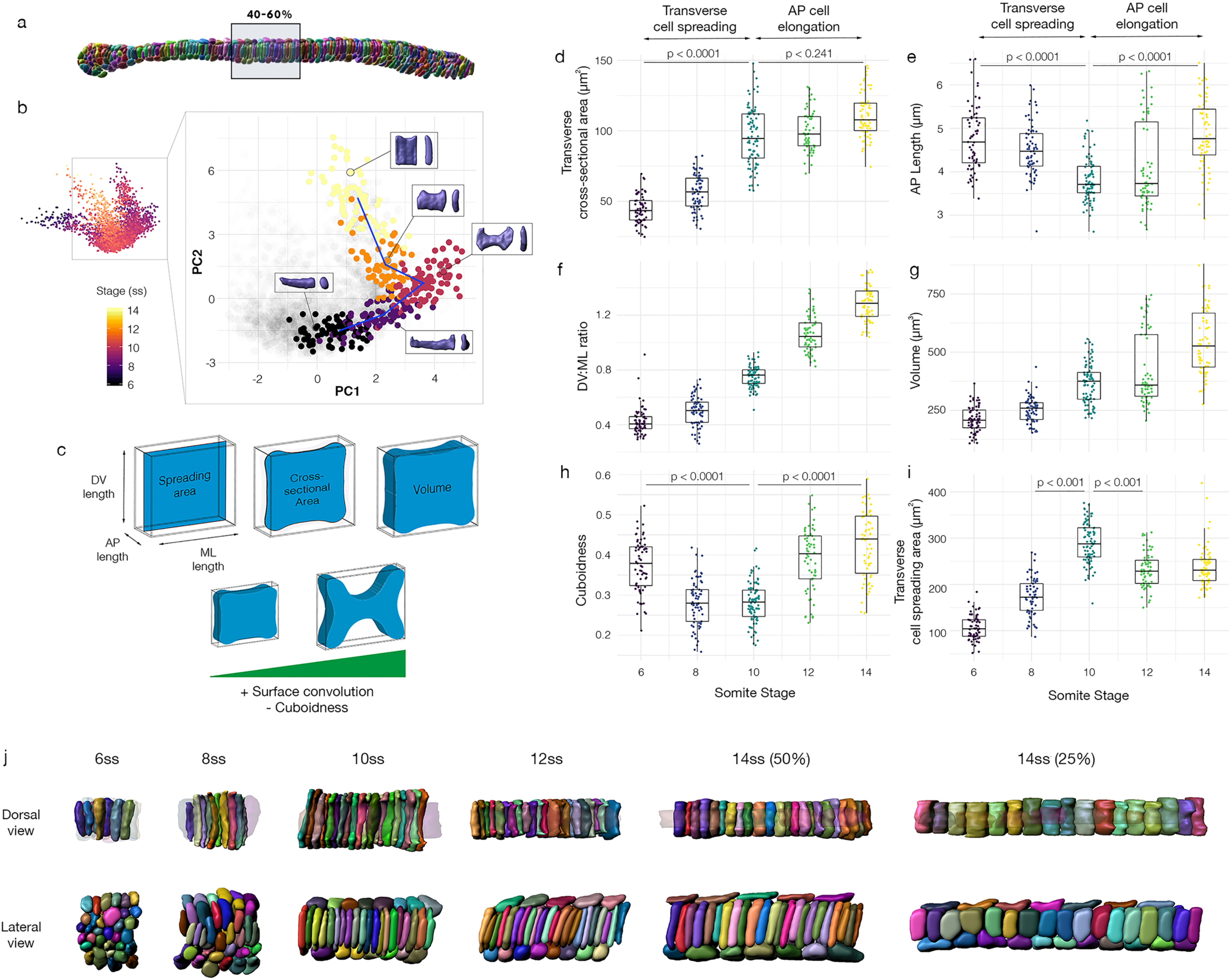
Cells in the central notochord layer pursue a two-step shape trajectory. (a) Representative 10ss notochord showing the region of cells sampled for the 50% level trajectory. (b) Trajectory reconstructed for central progenitors from the 50% level, shown in (a), with representative cells for each stage. (c) Shape parameterisation used to describe geometric transformations, based on cell volumes, bounding box dimensions and surface convolution values. (d – i) Quantification of individual geometric transformations; transverse cross-sectional area (d), AP length (e), DV:ML ratio (f), volume (g), cuboidness (h) and transverse spreading area (i). p values are described between groups using an unpaired two-tailed Student’s t-test. (j) Dorsal (top row) and lateral (bottom row) views of communities of 35 progenitors included in the trajectory in (a), showing temporal changes in topology, and an equivalent community from the 25% level, showing further elongation despite common topology and cell number.

To decompose these sequential transitions into their constituent geometric transformations, we analysed changes in the major correlates of each PC. Our shape parameterisation used for these descriptions is illustrated in *Fig. 2c* (detailed explanation in *Appendix I*). At 6ss, cells are clustered in morphospace based on a simple morphology that is elongated along the ML axis and rounded in lateral section (*Fig. 2a, 6ss*). In the first major shape transition (6ss – 10ss), progenitors undergo a decline in AP-oriented anisotropy (anti-correlated with PC1; *Fig. S1b, S3a*), such that transverse cross-sectional area increases at the expense of AP length (*Fig. 2d, e*). This means that the area of surface contact between adjacent cells increases. In parallel, the transverse profile of each cell is altered by sequential phases of elongation on ML and DV planes, that lead to a progressive increase in DV:ML ratio (positively correlated with PC2, *Fig. 2f*). At 8ss, cells reach their maximum elongation on the ML axis, with flattened tips at their lateral edges (*Fig. 2b, S3b*). By 10ss cells also spread extensively across the DV axis through a doubling of DV length (*Fig. 2b, S3c*). During the first transition, we also measured an increase in volume (positively correlated with PC3, *Fig. 2g*), and a significant decline in cell cuboidness, which reflects how faithfully the cell fits its object-oriented bounding box (anti-correlated with PC4, *Fig. 2h*). This means that the total reach of the cell is increased by convolution of its surface. Accordingly, we found the fall in cuboidness to coincide with a transient increase in cell spreading area (transverse bounding box cross-sectional area, *Fig. 2i*) to its maximum value at 10ss. Collectively, these data correspond to formation of a unique cell shape at 10ss, characterised by a medial point of constriction and flared lateral margins, similar in shape to a bowtie (*Fig. 2a, 10ss*).

During the second major transition (10ss – 14ss), we found cells to lose their bowtie morphology, and commence elongation along the AP axis (*Fig. 2a*). We identified an increase in cuboidness after the 10-somite stage (*Fig. 2h*), describing loss of the bowtie state and a return to more regular cuboidal morphologies (*Fig. 2b*). This also aligned with a decline in cell transverse spreading area (*Fig. 2i*). The loss of spreading area involved a specific retraction of ML cell length (*Fig. S3b*), whereas cells continued to elongate along the DV axis (*Fig. S3c*). As a result, the DV:ML ratio continued to increase during the second shape transition, thereby generating tall cuboidal cells (*Fig. 2f*). After 10ss, AP-oriented anisotropy now increased, corresponding to a decline on PC1 (*Fig. 2b*). In this period of AP elongation, we found the rate of increase in cell cross-sectional area to fall (*Fig. 2d*), and cell AP length to increase to match its value at 6ss (*Fig. 2e*). Given the ongoing increase in cross-sectional area between 10ss and 14ss (*Fig. 2d*), we reasoned that AP length must instead be increased through cell growth. Indeed, we found cell volume to increase at its highest rate during the second major transition, by a mean of 55% (*Fig. 2g*). These geometric transformations account for the emergence of tall cuboidal cells at 14ss (*Fig. 2b*). In sum, decomposition of the PCs revealed AP anisotropy, surface convolution and volumetric growth as the primary geometric transformations underlying the major shape transitions of the central cell trajectory.

The sequence of cell shape changes we identified in central notochord cells define a unique trajectory through morphospace. The next question is how these behaviours correspond to changes in multicellular organisation. To address this question, we segmented neighbourhoods of 35 adjacent cells at the 50% level, and aligned changes in their organisation with the transitions in cell shape defined above (*Fig. 2j*). At 6ss, all cells were elongated along their ML axes, with bipolar contacts with both the left and right margins of the chordamesoderm (*Fig. 2j, Dorsal view, 6ss*). Because this stage precedes most of notochord AP elongation (*Fig. 1a*), ML intercalation cannot generate further tissue length. Instead, we noted an increase in cell layers on the DV axis between 3ss and 6ss, suggesting that ML intercalation might drive an early convergent thickening (*Fig. S3 compare 3ss and 6ss, Fig. 2j Lateral view*). During the first shape transition in central cell progenitors (6ss – 10ss), in which they increase their transverse spread and adopt a bowtie morphology, we observed a process of intercalation oriented along the DV axis, that reduces the number of cell layers from 6 to 3 (*Fig. 2j, Lateral view, 6ss – 10ss*). During this DV intercalation, we found that cell neighbourhoods elongated, despite the AP shortening of individual cells (*Fig. 2j, 2e*). We found that DV intercalation was followed by a latter phase of neighbourhood elongation in the absence of further cell rearrangement, coincident with the second major transition in cell shape, in which cells increased in volume and AP length (*Fig. 2j 10 – 14ss, 2e, 2g*). This latter elongation phase was most pronounced in the pharyngeal region (*2j, compare 14ss 50% and 25*%). In sum, the sequential shape changes of the central cell trajectory aligned with distinct phases of notochord elongation, mediated first by DV cell intercalation, and second by cell elongation and growth.

### Single-cell morphometrics reveals temporal gradients in cell shape differentiation across the anteroposterior axis

We next applied our single cell morphometric approach to test for temporal variation between cells as they advance along the central cell trajectory. To achieve this, we used the spatial information associated with each set of cell shape descriptors to colour code cells in morphospace according to their position along the AP axis of the embryo (*Fig. 3a-d*). We grouped cells from different regions, either from the anterior 15% or posterior 85% of the notochord and analysed their distribution in morphospace at successive time points of development (*Fig. 3a-d*). By spatially mapping the data, we found that cells at different positions along the AP axis transit through morphospace at different times. Consistently, we observed that the most advanced cells along the central cell trajectory were located in the middle part of the notochord (*Fig. 3a-d, asterix marks leading edge in morphospace*). This initially consists of a single population of cells undergoing a common series of shape transitions (*Fig. 3a*). However, between the 12-somite and 14-somite stages, a distinction emerges between adjacent populations in the pharyngeal and trunk regions, which resolve on PC2 and PC3 (*compare Fig. 3cii with dii*). These mature states lie at the leading edge of a shape continuum that extends back to the most immature morphologies towards the posterior tip of the embryo (*Fig. 3a-d*). Indeed, in the posterior half of the notochord

**Figure 3.**
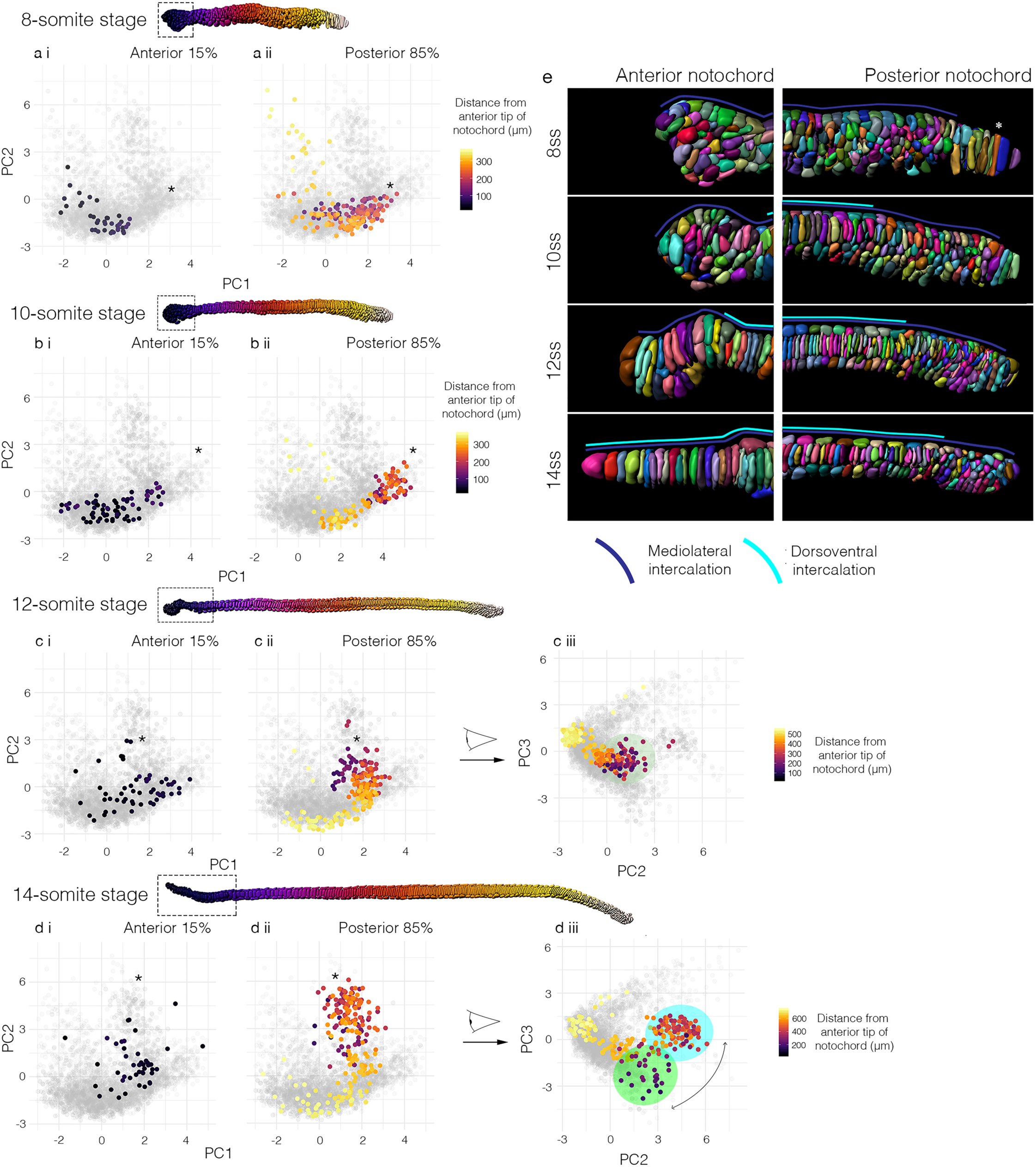
Morphospatial mapping of whole notochords reveals spatial variation in developmental timing. (a-d) Stack-of-coins progenitors from whole notochords at the 8- (a), 10- (b), 12- (c) and 14- (d) somite stages mapped into morphospace. Cells in the anterior 0.15 of axial length (boxes shown in each notochord) are mapped into a single graph (left, panel i), and the rest in a separate graph (right, panel ii). Colour-code reflects distance from the anterior tip of the notochord. c iii and d iii show the same cells dispersed on PC2 and PC3, revealing spatial separation of cells in either side of the bifurcation event in the stack-of-coins trajectory – cells in the pharyngeal region are qualitatively different to those in the more posterior trunk. (e) View of segmented anterior (left) and posterior (right) tips of the notochord at the 8-14 somite stages, showing temporal delays in mediolateral and dorsoventral intercalation relative to the central progenitors.

we find a strong positive correlation between the extent of shape differentiation and position along the AP axis (Pearson correlation coefficient with PC2 = 0.923, *Fig. S4c*). In addition, we identified a temporal lag in the development of the most anterior notochord progenitors. At the 10-somite stage, cells in the anterior 15% occupy the same position in morphospace as those at the 50% level do at the 8-somite stage (*compare Fig. 3ci and bii*). This asynchrony is also maintained at later stages, such that, at the 14-somite stage, anterior progenitors overlap with 50% progenitors at the 12-somite stage (*compare Fig. di and cii*); a consistent lag of approximately two somite periods. In sum, this analysis reveals bidirectional temporal gradients of cell shape differentiation, propagating from the centre of the notochord towards its anterior and posterior tips.

We now sought to test whether these temporal gradients of cell shape maturation were matched by corresponding AP differences in multicellular organisation. At all stages analysed, we found that progenitors in the anterior notochord exhibited delayed dorsoventral intercalation. Unlike the 50% level (*Fig. 2j*), they remained stratified on the dorsoventral axis until the 10-somite stage, and reduced to a single layered organisation by the 14-somite stage (*Fig. 3e, Anterior notochord, light blue line*). This represents a 4-somite delay compared to the trunk region (*Fig. 2j, 6ss – 10ss*). We additionally found the posterior notochord to exhibit delayed intercalation. At all stages analysed, progenitors in the most posterior notochord remained stratified on the dorsoventral axis (Fig. 3e, *Posterior*). Up to 12ss, we also found the persistence of persistence of tall columnar cells lacking bipolar left-right contacts at the extreme posterior tip of the notochord (*Fig. 3e, Posterior, asterix*). This is characteristic of the monolayered archenteron roof prior to mediolateral cell intercalation (*Fig. S5*). Over developmental time, cells organised in both of these patterns were progressively depleted and restricted further towards the posterior tip of the embryo, as sequential waves of mediolateral and dorsoventral intercalation spread across the tissue. This is also reflected in a gradual decline in occupancy of the central branch of the notochord morphospace between the 6-somite and 10-somite (*Fig. 1d, 3a-c*). In sum, this analysis shows that temporal gradients of cell shape differentiation across the AP axis are mirrored by corresponding delays in the timing of intercalation.

### Spatial variants in trajectory structure demonstrate divergent and convergent paths to specific cell morphologies

Having identified spatial variation in developmental timing across the AP axis, we sought to test whether all cells transition through the same sequence of shape changes irrespective of their spatial position. We investigated this by constructing region-specific shape trajectories for four regions sampled across the AP axis (*Fig. 4a*); anterior (0-15% AP position), pharynx (15-40% AP), trunk (40-60% AP) and posterior (60-100% AP). In each sampled region of the AP axis, we identified a unique variant of the central cell trajectory. In contrast to the trunk progenitors, we found anterior progenitors to be unique in their relative lack of structure in morphospace, with a significant overlap between cells from different stages (*Fig. 4bi*). Nonetheless, anterior cells did proceed along the same axes of variation as cells in the trunk region, albeit at a lesser magnitude (the 50% level). This included a transient loss of AP length during intercalation (*Fig. S6a*), and continuous increases in cross-sectional area (*Fig. S6d*), volume (*Fig. S6i*) and DV:ML aspect ratio (*Fig. S6j*). Maturation additionally involved a decline in the coefficient of variation for cuboidness, although the mean values remained constant (*Fig. S6g*), and a progressive decline in sphericity (*Fig. S6h*). This measures a process of stabilisation to more regular cell geometries over time through loss of surface convolution. Collectively, these patterns reveal similarity in the shape trajectory of anterior cells and those in more posterior regions, but suggest that there is a higher degree of local cell shape variation as cell progress to their final morphologies (*Fig. 4b*).

**Figure 4.**
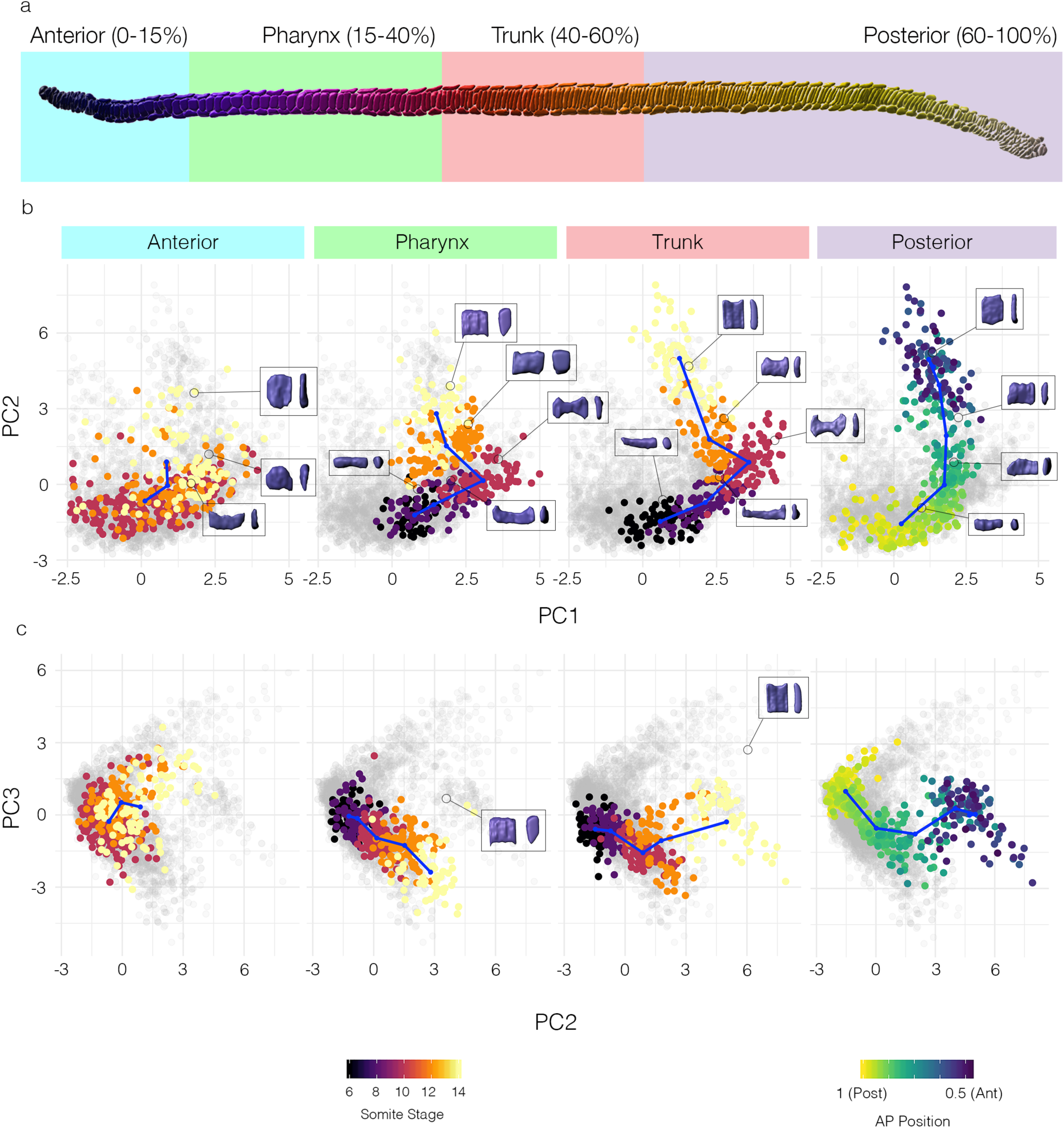
The trajectories of central notochord cells exhibit predictable spatial variation. (a) 14ss notochord highlighting each region sampled for trajectory construction. (bi - ei) PC1 vs PC2 for each region of the AP axis with colour-code for developmental stage (anterior, pharynx, trunk) or anteroposterior position (posterior). Inlays show representative cells for each region marked in the morphospace. (bii - eii) PC2 vs PC3 for each AP region shown. Inlays illustrate different morphologies emerging from the central cell trajectory in the pharynx and trunk.

The trajectories followed by cells at the pharyngeal and trunk levels are similar to one another in that they display the two major transitions observed at the 50% level, including entry to the ‘bowtie’ domain at the 10-somite stage (*Fig. 4ci, di*). In both cases, and unlike in the anterior notochord (*Fig. 4bi*), cells transitioned collectively during development, as illustrated by non-overlapping occupancy of morphospace at successive stages (*Fig. 4ci, di*). However, a major difference in the pharyngeal population, compared to the trunk, was greater and more rapid changes in length and volume between the 12- and 14-somite stages (*compare Fig. 2e, g with Fig S7a, i*). Pharyngeal cells also spread less extensively across PC2 because they stabilise a square transverse profile (mean DV:ML ratio of 1.035), rather than elongating along the DV axis in the second major shape transition like in the trunk (mean DV:ML ratio of 1.283) (*compare Fig. 2c and Fig 7j*). As a result, the central cell trajectory exhibits a second bifurcation event in morphospace across PC2 and PC3, yielding two unique cell morphologies that are spatially resolved along the AP axis in the pharynx and trunk (compare *Fig. 4cii, dii*). Collectively, in this analysis we find that notochord cells at the pharyngeal and trunk level initially follow a common trajectory that branches to divergent terminal morphologies.

To predict the trajectory of remaining posterior progenitors (the posterior 40% notochord length), which continue to differentiate beyond the final timepoint of analysis, we used relative AP position to organise them in a pseudotemporal order. This was justified by the strong correspondence we identified previously between axial position and developmental maturity in this posterior region (*Fig. 3a – d, Posterior panels*). For the posterior trajectory, each landmark was defined as the mean coordinate position for cells occupying 5 evenly-sized bins of the AP axis, from posterior to anterior. In this region, we found that cells appeared to totally circumvent a bowtie morphology, as shown by failure to populate the ‘bowtie’ domain occupied by pharyngeal and trunk cells at 10ss (*Fig. 4ei*). This implies a shortcut in their trajectories, such that maturation involves continuous elevation of transverse spreading area (increase in on PC1), with little or no surface convolution. In sum, this analysis reveals discontinuity in shape transitions across the AP axis, in which some trajectories split over time to generate morphological diversity, while others take divergent paths to common morphologies.

### Geometric modelling reveals a requirement for growth in coupling convergence and extension

By deconstructing notochord development in morphospace, we have shown that AP anisotropic elongation, growth and surface convolution are key geometric transformations underpinning central cell differentiation (Fig. 2). Therefore, changes in cell length should be the summed effect of AP anisotropic elongation, which we know has an adverse effect on cell length, and growth acting together. We next set out to investigate the effect of each transformation on the lengths of single cells, and groups of cells undergoing intercalation, when applied individually. To this end, we devised a simplified *in silico* framework to model the effect of each transformation on a mean cell from the 6-somite stage notochord at the 50% level. This cell is defined by its AP length 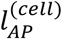, volume *V*^(*cell*)^, and cross-sectional area *A*^(*cell*)^. Where *s* denotes stage, AP length can be calculated as

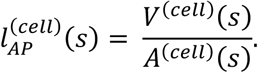

We can now measure change in cell length when change in either volume or cross-sectional area occur independently, with the other fixed at their 6ss value.

As we have shown experimentally, central notochord cells in the 50% region undergo an early phase of shortening on the AP axis linked to intercalation (6-10 somites), followed by a late phase of elongation in which AP length is restored (10-14 somites) (*Fig. 2e, 5b black line*). We first set out to investigate how cell length would change if governed only by AP anisotropy, in the absence of growth. Here, we allowed cell length to change in accordance with measured change in cross-sectional area, while maintaining the cell at constant volume (Fig. 5b, blue line). Under these modelled conditions, cross-sectional area increases at the expense of length, such that cells undergo a 2.16-fold shortening between 6ss and 10ss. Thereafter, instead of restoring length, cells continue to shorten at a slower rate, by an additional 1.17-fold change. This suggests that, during intercalation, transverse cell spreading has a sustained negative effect on length. Without prior knowledge of growth dynamics, we envisaged two scenarios for its effect on shape. First, we considered a scenario in which cells maintain their shape at the 6ss, and the growth that we measure (*Fig. 2g*) acts isotropically to expand cell geometry proportionally in all directions (*Fig. 5b, grey line*). In this case, cells underwent a progressive increase in length by a total of 1.37-fold, thereby exceeding the 1.02-fold net length change measured during normal development (*Fig. 5b, grey line*). This change was more pronounced in the second scenario of anisotropic growth, in which we prevented radial cell expansion and forced growth to act unidirectionally on AP length. In this case, we measured a 2.59-fold elongation (*Fig. 5b, orange line*), again far exceeding the measured change. Collectively, these calculations suggest that transverse cell spreading behaviour and growth have antagonistic contributions to cell length, and the real profile of length change is a dynamic balance between the two transformations. Up to the 10-somite stage, the rate of transverse cell spreading outweighs that of growth, leading to a net cell shortening during intercalation. When cross-sectional area stabilises at 10ss, growth then dominates to translate a programme of transverse cell spreading into one of elongation.

**Figure 5.**
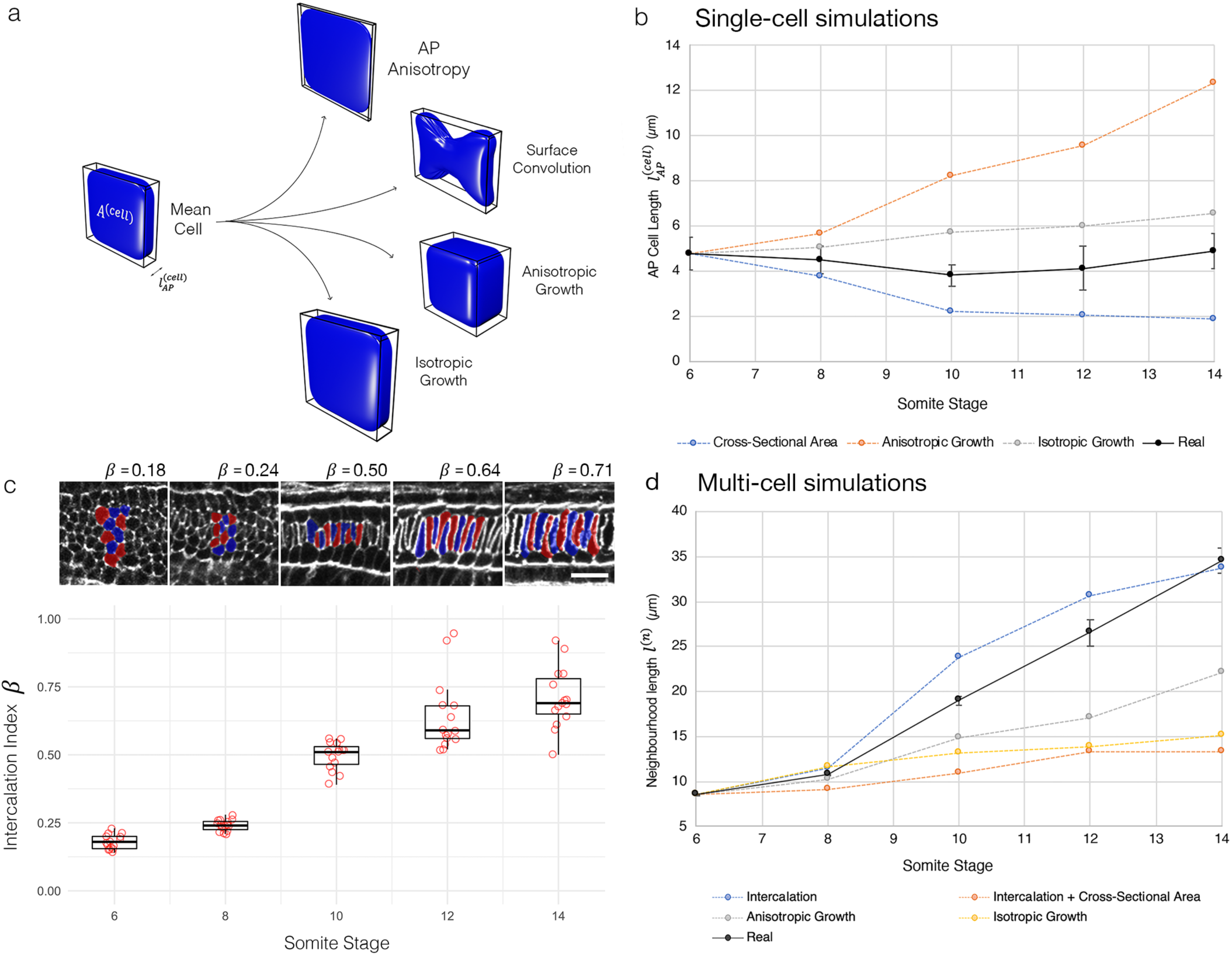
Geometric modelling predicts contributions of shape and size to major transitions of the central cell trajectory. (a) Visual renderings of transformations identified in the central cell trajectory. (b) Results of geometric perturbations applied to a mean central cell from the 6-somite stage, over a developmental time course, illustrating predicted change in cell length. Coloured lines represent modelled scenarios, black line represents real measured change with measured standard error. (c) Quantification of intercalation index across developmental time, with representative groups of 10 cells per stage. Scale bar shows 20µm. (d) Predicted change in the length of a neighbourhood of 10 6-somite central cells, integrating both intercalation and changes in single-cell shape and size. Black line shows real measured length changes, with standard error.

We next sought to test how this relationship between shape and size in single cells affects the rate of elongation in a group of neighbouring cells undergoing intercalation. We tested this in groups of 10 cells from the 50% region of the notochord. To investigate the effect of intercalation, we defined an intercalation factor, *β*, which allows calculation of extended AP group length *l*^(*n*)^ from cell number *n* and mean AP cell length 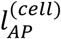 for a given stage (*Fig. 5c, appendix I*);

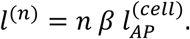

We made use of values for *l*^(*n*)^ obtained from groups of 10 adjacent cells measured in the embryo at each stage. When *β* = 1, group length equates to the summed lengths of all cells, whereas when *β* < 1 cells are displaced from the midline and so their individual lengths are not additive in the same plane (*Fig. 5c, Fig. A2*). We used these metrics to drive intercalation either independently, or in combination with transformations in cell shape and size. We first tested how intercalation alone contributes to length by applying experimentally obtained intercalation index values (*β* values) (Fig. 5c) to groups of mean 6ss progenitors. Here, we found that intercalation can drive a 3.49-fold AP elongation, occurring at its greatest rate between the 8-somite and 12-somite stages (Fig. 5d, blue line). This is the effect of simply aligning cells into a single-file array, with no accompanied shape change. To then factor in cell shape changes, we applied measured changes in both cross-sectional area (as for Fig. 5b) and intercalation index, while keeping volume constant (Fig. 5d, orange line). In this growth-free scenario, intercalation did increase neighbourhood length, despite the shortening of individual cells, but only to a maximum 1.55-fold extension at the 12ss. This falls significantly short of real measured values, therefore suggesting that the ability of cell intercalation to drive AP tissue elongation is counteracted by cell spreading behaviours that reduce cell length. We therefore hypothesised that growth should account for most of AP elongation. By itself, isometric growth achieved only a 1.76-fold elongation (Fig. 5d, yellow line), whereas anisotropic growth was more effective, driving a 2.59-fold increase (Fig. 5d, grey line). This means that growth is also insufficient to drive full tissue elongation, unless coupled to intercalation. In sum, our geometric modelling suggests that growth enables convergent extension in two manners. First, by counteracting loss of cell AP length during intercalation due to cell spreading (6ss – 10ss). Second, by further increasing cell length after intercalation, once the stack-of-coins is formed (10ss – 14ss).

### Posterior axial progenitors accelerate notochord elongation by increasing cell number

Our approach of mathematically deconstructing and reconstructing the central cell trajectory predicted a balance of shape regulation and growth that results in a constant rate of elongation in cell neighbourhoods from 8ss onwards. Our final objective was to test whether the behaviours we identify at this scale are sufficient to account shape change at the tissue-scale. We therefore compared the dynamics of elongation at the neighbourhood level with that at the tissue level, using direct measurements of total notochord length (*Fig. 6a*). Here, we found the amount of whole notochord elongation to exceed that of local cell neighbourhoods, due to a specific acceleration in rate between 8ss and 10ss (*Fig. 6b*). This discrepancy indicated that some other factors apart from change in cell shape and size must be affecting the rate of tissue elongation. We therefore hypothesised that addition of new cells through cell division might be required to explain full notochord elongation at the tissue-scale.

**Figure 6.**
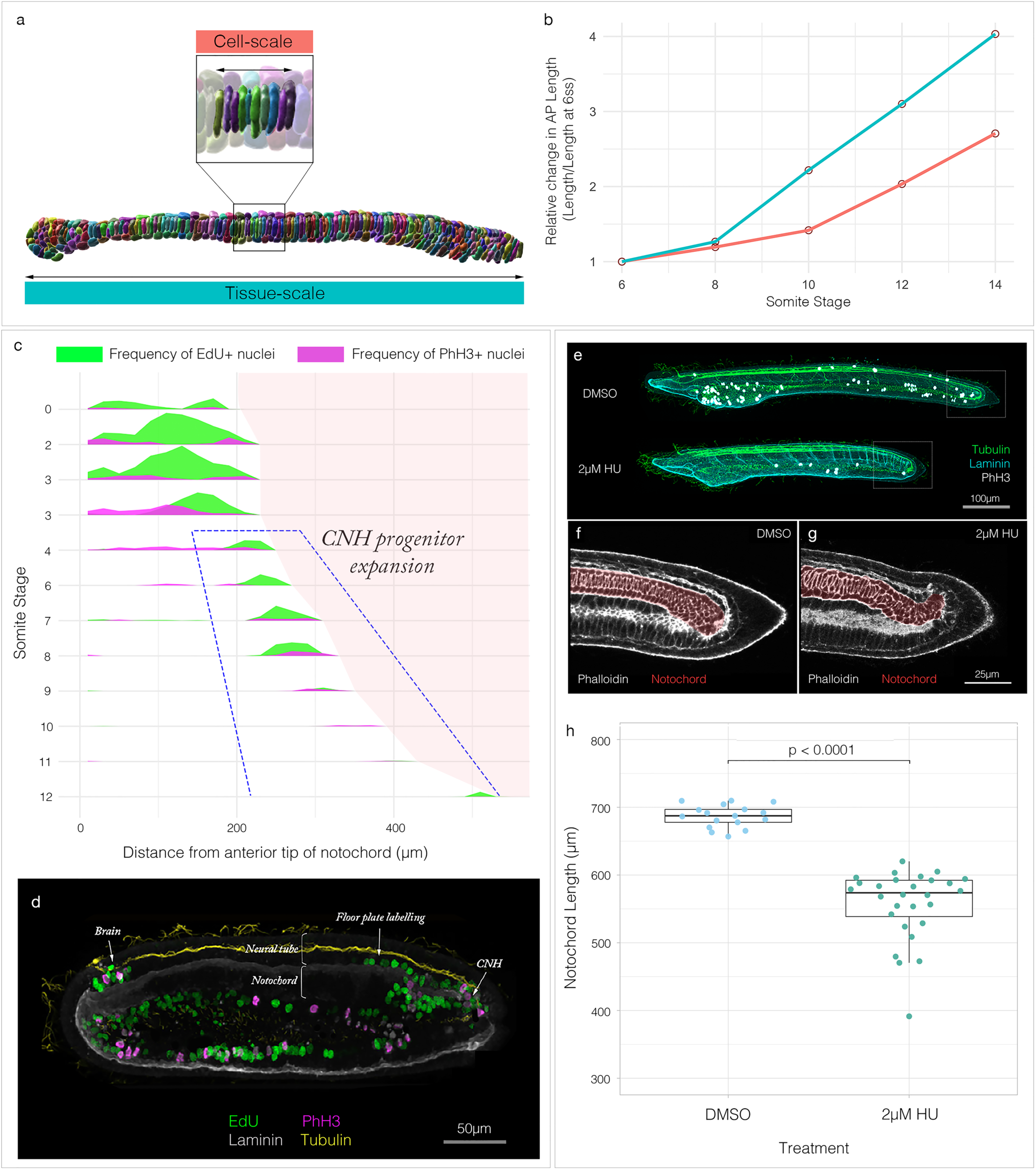
Posterior addition is required for full extension of the body axis. (a) 10ss segmented notochord illustrating cell-scale (top) and tissue-scale (bottom) measurements of AP length. (b) Graph showing relative change in AP length at the cell-scale (neighbourhoods of 10 adjacent cells) and tissue-scale (whole notochords). (c) Tissue-specific proliferation landscape for the notochord, showing mean frequency of EdU and PhH3 staining in embryos at successive somite stages, with length scaled to mean per stage (*n =* 102). (d) Raw staining for EdU and PhH3 at the 8-somite stage in a sagittal section. During elongation, proliferation restricts to the anterior neural plate, and chordoneural hinge in dorsal tissues, and continues in the ventral endoderm. Staining around the chordoneural hinge (CNH) extends into the posterior floor plate and notochord. (d) Effect of 2µM hydroxyurea on axial length from the 6-somite stage. (f, g) Phalloidin staining in a sagittal section through tails of DMSO and HU-treated specimens, notochord highlighted in red. (h) Quantification of total axial length in DMSO and HU-treated specimens (*n* = 17 DMSO, 27 HU).

To test our hypothesis that cell division modifies the notochord elongation curve by increasing cell number, we first set out generate a map of cell division dynamics in the notochord during its elongation. With this objective, we first labelled embryos at successive somite stages with markers for nuclei in two cell cycle phases; EdU, to cumulatively label cells passing through S-phase in the time window of exposure (applied 2-hours before fixation), and immunostaining for phosphorylated histone 3 (PhH3), to mark mitotic cells at the moment of tissue fixation (*Fig. S8*). We then plotted the frequencies of labelled cells across developmental time and space to construct a tissue-specific proliferation landscape (*Fig. 6c*). This landscape illustrates a dynamic pattern of proliferative domains in the notochord, which become sequentially enriched with EdU and PHH3, thereby capturing cell cycle progression. Prior to elongation, these data reveal broad cell division throughout the notochord (*Fig. 7c, 0 – 4ss*). However, at the onset of elongation (*Fig. 6c, 6ss*), this transitions to a specific proliferative domain at the posterior tip, which is active until the 8-somite stage (*Fig. 6c 8ss, 6d*). Therefore, we can infer that cells developing along shape trajectories defined for the anterior, pharyngeal and trunk regions of the notochord are in a post-mitotic state during tissue elongation (not labelled with EdU or PhH3, *Fig. 6c*), whereas the posterior trajectory is continually fuelled by cell division. To functionally test the contribution of posterior proliferative progenitors to tissue elongation, we treated embryos with hydroxyurea (HU) at 6ss, when proliferation becomes specifically restricted to the posterior tip. We observed that HU-treated embryos elongated, but to only 81% of their expected length by the 14-somite stage (*Fig. 6e - h*), and this was coupled to disorganisation of the posterior notochord, which lost its regular stack-of-coins pattern (*Fig. 6f, g*). In sum, while notochord length is primarily generated by cell rearrangement and growth, these data reveal a further role for posterior notochord progenitors in providing additional cellular material to accelerate tissue elongation.

## DISCUSSION

Here, we decompose the development of an entire tissue - the amphioxus notochord - by embedding it in a single-cell morphospace. In this environment, morphogenesis is unravelled into a branching portrait of shape differentiation, in which cells transit along trajectories specific to cell type to form a stereotypical diversity of morphologies. When spatially mapped in the embryo, these trajectories organise into temporal gradients of shape differentiation, which in turn align with stepwise changes in multicellular topology. By carrying spatial coordinates into the morphospace, our approach exposes both predictable variation in cell morphology across the AP axis, and the convergence of cells on common morphologies through variable morphogenetic routes. Furthermore, because our approach allows extraction of dynamic information from static imaging data, it is widely applicable in a range of organisms where genetics and live imaging are challenging. Our findings suggest that cell morphology may contain a much richer body of information than is typically assumed, to the point of being predictive of cell identity, spatial position, and developmental time. Single-cell morphometrics therefore stands to complement single-cell genomics as a rich resource for building multiscale definitions of cell types, and defining their roles in the emergence of embryonic form (Briggs et al., 2018; Ibarra-Soria et al., 2018; Sebé-Pedrós et al., 2018; Wagner et al., 2018). In this case study, we have uncovered remarkable complexity in the trajectories of cell shape change responsible for amphioxus notochord formation, offering a unique window into emergence of a morphological novelty at the base of the chordate phylum. Further studies will take advantage of developments in automated image segmentation and multivariate shape quantification to dissect morphogenesis in a diversity of systems, and integrate these data to define patterns of morphogenetic variation over both developmental and evolutionary time scales. In this respect, single-cell morphometrics will offer a quantitative framework for comparative morphogenesis.

An unexpected finding in our study was regional variation of cellular behaviour across the AP axis, within a tissue that appears, at face value, to be morphologically continuous: In the anterior we find a highly variable morphogenesis, perhaps characterised by asynchronous maturation between neighbours; in the pharynx and trunk we find a unique dynamic in which a bowtie morphology disproportionately expands cell spreading area during intercalation; and in the posterior we find a simplified shape trajectory fuelled by cell division. These variations may arise due to a common programme of differentiation occurring under unique mechanical conditions or be the effect of region-specific signalling and genetic regulation. Such correlation is supported by the explicit notochord regionalisation found across vertebrates. In the mouse, live imaging has revealed marked differences in morphogenesis between the anterior head process (AHP), trunk and tail notochord (Yamanaka et al., 2007). Unlike the trunk, AHP progenitors derive from the early- and mid-gastrula organiser without passing through the node (Kinder et al., 2001; Yamanaka et al., 2007), and accordingly are uniquely sensitive to loss of Nodal signalling, but are insensitive to loss of Noto expression (Vincent et al., 2003). In contrast, the tail progenitors are unique in their active migration to move posteriorly to the node late in axis elongation (Yamanaka et al., 2007). Another example of regionalisation is the prechordal plate (PrCP) in all vertebrates, which shares with the AHP its developmental origin from the most anterior axial mesoderm, and forms through anteriorly-directed collective cell migration (Tada and Heisenberg, 2012). Evidence for regionalised behaviour aligns with nested expression of Hox genes in the vertebrate notochord (Prince et al., 1998). Previous work has also identified regionalised gene expression in the amphioxus notochord, and the data presented here adds to this in hinting at readout of these variations as discrete cell behaviours (Albuixech-Crespo et al., 2017). In evolution, the striking variations in vertebrates – the PrCP, AHP, trunk and tail regions – may have emerged within semi-discrete morphogenetic fields that were already defined in the first chordates.

By deconstructing and reconstructing the trajectory for central notochord cells, we elucidate a balance of cell shape, size and topology that dictates the length of small cell neighbourhoods. First, we infer that mediolateral intercalation does not contribute directly to notochord elongation. Rather, it drives a convergent thickening, that increases the number of cell layers stratified across the dorsoventral axis. This multi-layered organisation is then reduced to a trilaminar pattern through dorsoventral intercalation, which is linked to convergent extension and tissue elongation at the tissue level. In this second process, we find that individual notochord cells increase their transverse spreading area at the expense of their AP length. While transverse cell spreading may facilitate intercalation between neighbouring cells, our geometric modelling suggests that the coupled loss of cell length almost entirely abrogates the contribution of intercalation to tissue elongation. In turn, we find that cell growth is required to counterbalance cell length. As a result, we predict that cell growth is required in this system to ensure tissue elongation. During intercalation, cell growth buffers the loss of cell AP length due to cell spreading, thereby enabling cell intercalation to generate tissue length. This accounts for the first phase of notochord elongation. After intercalation, cell growth plays an additional role in further increasing cell length, once cross-sectional area is stabilised. This enables tissue length to further increase, without any ongoing changes in cell topology, at a constant rate. At this scale, we therefore predict that a tight spatiotemporal coordination of growth is required in the notochord to modulate cell shape and in turn control tissue shape and size. Conversely, the effect of growth on form is equally controlled by active changes in cell shape and topology.

Our investigation also revealed an important role for cell division in posterior axial progenitors for defining notochord length at full extension. Indeed, neighbourhood elongation dynamics alone are insufficient to account for those at the tissue scale. We show that this discrepancy may be explained by cell division during notochord elongation, which generates cells for its posterior 20%. It is important to note that the programme of cell shape morphogenesis that generates length operates only after cell division arrest, when waves of intercalation and growth propagate across the tissue. As such, cell division, intercalation and growth are temporally separated, but ultimately act cumulatively to generate tissue length. Our data suggests that cell division is not necessarily a length-generating process, rather it dictates the number of cells available to fuel length-generating mechanisms that act later in development. This stresses the importance of studying morphogenesis over broad developmental time scales. The amount of cell division we find in amphioxus, and its contribution to length, is relatively small compared to vertebrate systems like mouse and chick (Bénazéraf et al., 2017; Steventon et al., 2016). However, its presence in amphioxus is important in offering an evolvable node for evolutionary change. This is supported by an increase in the role of cell division in the notochord elongation throughout vertebrate evolution, contributing to an increase in its size and length. In amniotes, extensive cell division and growth in posterior progenitors dramatically expand the size of the notochord field when it is established during gastrulation (Catala et al., 1996; Mugele et al., 2018; Selleck and Stern, 1991). Our data suggest that these dynamics are not inherently novel, rather they have arisen through changes in the magnitude of cell division in cell types already present in the first chordates.

The approaches presented here offer a new way of seeing in the study of tissue morphogenesis, that enables holistic analysis of cell behaviours defining tissue geometry and lends itself to cross-species comparisons. In this case study, we use single-cell morphometrics to define a new model for notochord morphogenesis in the amphioxus, and in doing so shine light on principles of morphogenesis at the base of the chordate phylum. We find a conserved role for cell intercalation and growth in generating notochord length that complements previous studies in vertebrates, and also identify a number of evolvable nodes that predict the diversity of developmental dynamics found in vertebrate model systems. This includes evidence for spatial variation in developmental timing across the anteroposterior axis, localised differences in single-cell shape trajectories within a morphological continuum, and a role for axial progenitor cells in accelerating notochord elongation through cell division. As a result, we propose that the diversification of notochord form and developmental dynamics in vertebrates has depended more on tweaks in the magnitude of morphogenetic processes already present in the first chordates, rather than their innovation *de novo*.

## MATERIALS AND METHODS

### Animal husbandry, spawning and fixation

Wild catch collections of amphioxus, *B. lanceolatum*, were made in Banyuls-sur-Mer, France, and transported to a custom-made amphioxus facility in Cambridge, UK. Adult amphioxus were maintained, bred and the progeny raised as described in Benito-Gutierrez et al (2013). All embryos were fixed in 3.7% PFA + MOPS buffer for 12 hours, then stored in sodium phosphate buffered saline (PBS) + 0.1% sodium azide at 4°C.

### Embryo staining and imaging

Embryos were first permeabilised overnight in PBS + 1% DMSO + 1% Triton. They were then blocked in PBS + 0.1% Triton + 0.1% BSA + 5% NGS, and incubated overnight in primary antibodies as follows: rabbit anti-laminin (Sigma, L9393) at 1:50, rabbit anti-PhH3(Abcam, ab5176) at 1:500, mouse anti-acetylated tubulin(T 6793, Sigma) at 1:250.

Primary wash was performed in PBS + 0.1% Triton + 0.1% BSA, before a secondary block in PBS + 0.1% Triton + 0.1% BSA + 5% NGS, and overnight incubation in goat anti-rabbit and/or goat anti-mouse secondary antibodies at 1:250. Staining with DAPI at 1:500 and rhodamine phalloidin at 1:250 was performed with the secondary incubation. Embryos were washed thoroughly with PBS + 0.1% Triton and mounted for confocal imaging on glass-bottomed dishes in 80% glycerol. All imaging was performed on an Olympus V3000 inverted confocal microscope at 30X optical magnification.

For EdU labelling, EdU was applied to live embryos in seawater at a final concentration of 20µM for 2 hours prior to fixation. Fluorescent detection of incorporated EdU was performed following the manufacturer’s instructions using a Click-it EdU Alexa Fluor 647 Imaging Kit (Invitrogen) prior to primary antibody incubation.

### Drug treatment

Live amphioxus embryos were treated with 2µM hydroxyurea (Sigma, H8627) or dimethylsulfoxide (DMSO; Sigma, 276855) continuously between the 6-somite and 14-somite stages (18-34hpf at 21°C). They were then fixed immediately for imaging.

### Image segmentation and shape quantification

Z-stacks of embryos immunostained for tubulin and laminin, and stained for actin with phalloidin, were imported to Imaris 9.2.1 for segmentation.

Cells were segmented manually, using phalloidin staining to delineate cell outlines. Splines were drawn around each cell every 2 slices in parasagittal section. Cell surfaces were validated by false colouring the contents and checking its faithful fit to the phalloidin stain. Cells were excluded if contours could not confidently be drawn using the phalloidin stain. Most shape metrics were obtained directly from Imaris (raw xyz position for centre of surface homogenous mass, 3D orientation of the long ellipsoid axis, area, axis-aligned/object-oriented bounding box dimensions, centre of homogenous mass for the DAPI channel masked within each surface, number of triangles, volume). The remainder were calculated manually as stated, where *l* represents length, *x, y*, and *z* represent positions in space, *bb* refers to bounding box-derived metrics, and *cell* refers to cell-derived metrics. Position along the AP axis was also maintained for all cells.

| Quantity | Calculation |
| --- | --- |
| AP:DV ratio | $l_{AP}^{(bb)} / l_{DV}^{(bb)}$ |
| AP:ML ratio | $l_{AP}^{(bb)} / l_{ML}^{(bb)}$ |
| DV:ML ratio | $l_{DV}^{(bb)} / l_{ML}^{(bb)}$ |
| Mean cell diameter<br>$d^{(cell)}$ | $\frac{l_{DV}^{(bb)} + l_{ML}^{(bb)}}{2}$ |
| AP anisotropy | $l_{AP}^{(bb)} / d^{(cell)}$ |
| Transverse spreading area | $l_{DV}^{(bb)} \times l_{ML}^{(bb)}$ |
| Cross-sectional area<br>$A^{(cell)}$ | $\frac{V^{(cell)}}{l_{AP}^{(bb)}}$ |
| Cuboidness | $Vol / (l_{AP}^{(cell)} \times l_{DV}^{(cell)} \times l_{ML}^{(cell)})$ |
| Sphericity | $\frac{\pi^{\frac{1}{3}} (6V^{(cell)})^{\frac{2}{3}}}{A^{(cell)}}$ |
| Flatness | $l_{max}^{(cell)} / l_{min}^{(cell)}$ |
| Nuclear displacement from centre of<br>homogenous mass<br>(x, y, z) | $ABS(x^{(cell)} - x^{(DAPI)})$<br>$ABS(y^{(cell)} - y^{(DAPI)})$<br>$ABS(z^{(cell)} - z^{(DAPI)})$ |
| Transverse surface convolution | $\frac{A^{(cell)}}{l_{DV}^{(cell)} \times l_{ML}^{(cell)}}$ |

### Principal component analysis

Principal component analysis (PCA) was performed using the Factoextra package in R on the following geometric variables;

- Angle of major ellipsoid axis relative to x, y, z coordinate system
- Number of triangles
- Volume
- Relative AP, DV, ML dimensions
- AP anisotropy
- Nuclear displacement from centre of homogenous mass on x, y and z planes
- Cuboidness
- Flatness
- Compactness

The ‘scale’ function was used to standardise the data across all cells, giving them a standard deviation of 1 and mean of 0. The PCA coordinates were merged with the raw dataset to allow colour-coding and filtering for specific groups. For stage, we filtered based on the number of somites counted along the AP axis using phalloidin, and laminin immunostaining. For region on the AP axis, we sampled four broad domains; the anterior (0-15% axial length), pharynx (15-40% axial length), trunk (40-60% axial length) and posterior (60-100% axial length), where 0% is the anterior tip of the notochord and 100% is the posterior tip. For cell layer on the DV axis, the prospective Müller cells were filtered out as the most dorsal and ventral rows of notochord cells at each stage, regardless of morphology. All remaining cells were classified as ‘central’ cells. Morphospace graphs were constructed using *ggplot2*.

Morphospatial trajectories were constructed by connecting the centre point for point clouds at successive developmental stages, calculated as the mean position on each PC shown in the graph. For the posterior 40% of the notochord, which continues differentiating at 14ss, the trajectory was constructed by connecting the centre points for point clouds from five evenly-sized bins of the AP axis, from posterior to anterior. Here, relative AP position is used as a proxy for developmental maturity

### Cell position mapping

To construct proliferation landscapes, the point selection tool in FIJI/ImageJ was used to identify a coordinate position for each nucleus positive for EdU or PHH3, and the most anterior and posterior nuclei in the embryo. The position for each labelled nucleus was then quantified across a normalised AP axis. Data was then pooled for embryos at each somite stage, and normalised length was scaled to the mean per stage. Mean frequency of EdU+ and PHH3+ nuclei was then plotted using the ggridges package in R to generate proliferation landscapes.

## Acknowledgements

We thank Michael Akam, Andrew Gillis, John Marioni, Berta Verd and the EBGLab for critical reading of the manuscript and feedback. We also thank Matt Wayland in the Imaging Facilities at the Department of Zoology and the Cambridge Advanced Imaging Centre for their support and assistance.

## APPENDIX I: *IN SILICO* GEOMETRIC MODELLING

### Geometric modelling of single-cell shape changes

The aim of the *in silico* geometric modelling is to identify how different geometric transformations change cell and tissue length. We first need to define a set of measures that allow for a simplified characterisation of the cell shapes and shape changes. To this aim, we first create the object-oriented bounding box of the cell corresponding to the three-dimensional cell shape and find that the box aligns to the anterior-posterior axis (AP-axis), the dorsal-ventral axis (DV-axis) and the medial-lateral axis (ML-axis) of the embryo (see Fig. 1a). The bounding box has the volume *V*^(*bb*)^ and is defined by three lengths, 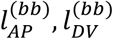 and 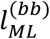 (see Fig. 1b). We assume that the cells can be approximated as a two-dimensional shape that is oriented along the *DV* − *ML* plane and then projected along the *AP* axis for a length 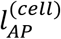 (see Fig. 1b and 1c). We find that bounding box length 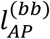 is a good approximation for cell length 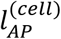, but convolution of the membrane on the transverse *DV* − *ML* plane generates a discrepancy between cell spreading area, defined by the bounding box *A*^(*bb*)^, and real cell area *A*^(*bb*)^. In this case, we can assume

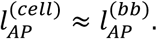

We then define the spreading area

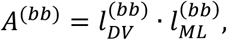

which corresponds to the transverse area (the area in the *DV* − *ML* plane) of the bounding box and the transverse area of the cell,

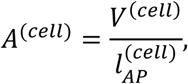

where *V*^(*cell*)^ is the cell volume. We also define the ratio

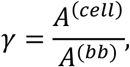

which is a measure of how convoluted the transverse cell shape is in the *DV* − *ML* plane. For a small *γ*, the cell is characterised by one more several long and thin elongations, for *γ* = 1 it fills the complete rectangle (see Fig. 1c). We now start with a cell at somite stage *s ∈* {6,8,10,12,14} which has the volume *V*^(*cell*)^(*s*), the anterior-posterior length 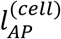 (*s*) and the ratio *γ*(*s*). In this case, the spreading area is given by

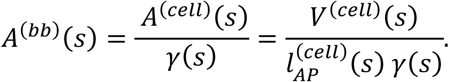

Alternatively, to calculate change in length 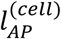, we start with a cell of volume *V*^(*cell*)^(*s*), area *A*^(*cell*)^ and ratio *γ*(*s*). The length 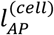 is now given by

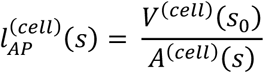

where *s*_0_ defines the initial state of the cell at stage *s*_0_. We find that the cell length along the *AP* axis only changes when the cell is elongated in this direction, grows anisotropically in this direction or isotropically in all directions. A convolution that does not affect the transverse area *A*^(*cell*)^(*s*) will also not affect the elongation.

**Figure A1.**
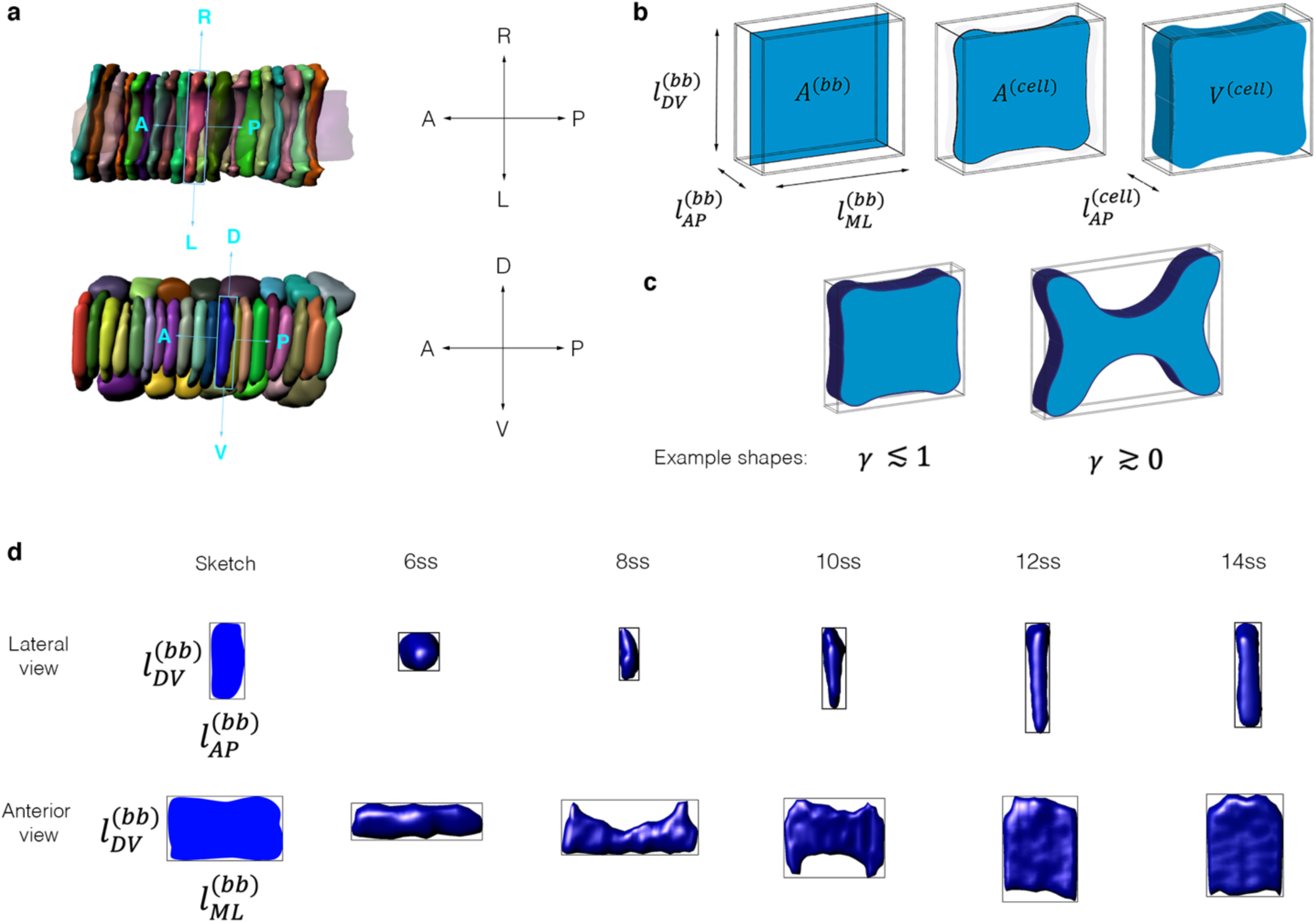
Cell shape metrics used for geometric modelling. (a) Object-oriented bounding boxes are calculated for each cell, and their axes are aligned to the AP, DV and ML axes of the embryo. (b) Schematised cells within object-oriented bounding boxes, showing the measurements of length, area and volume acquired. (c) Cells define the dimensions of their bounding boxes through variable degrees of surface convolution, which we quantify as *γ*. (d) Sample cells for each stage of development in lateral (top row) and anterior (bottom row) view within object-oriented

We can now study different types of geometric transformations and check how they affect these quantities:

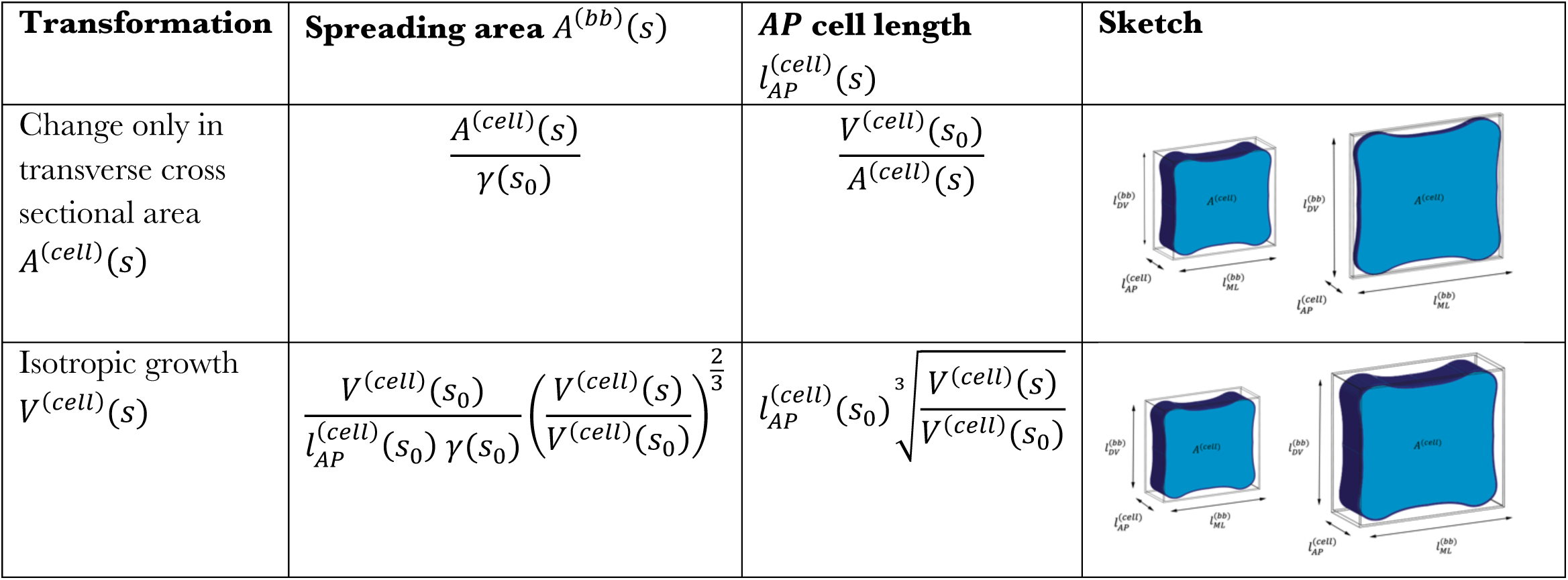
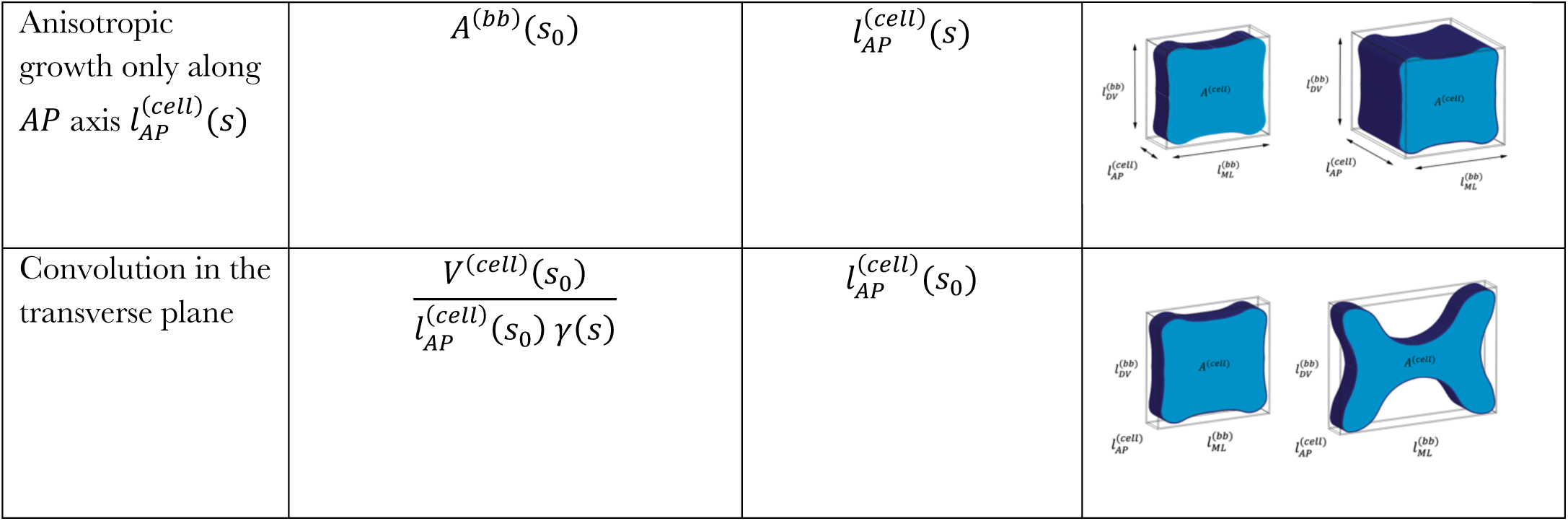

| Transformation | Spreading area $A^{(bb)}(s)$ | AP cell length $l_{AP}^{(cell)}(s)$ | Sketch |
| --- | --- | --- | --- |
| Change only in transverse cross sectional area $A^{(cell)}(s)$ | $\frac{A^{(cell)}(s)}{\gamma(s_0)}$ | $\frac{V^{(cell)}(s_0)}{A^{(cell)}(s)}$ | |
| Isotropic growth $V^{(cell)}(s)$ | $\frac{V^{(cell)}(s_0)}{l_{AP}^{(cell)}(s_0) \gamma(s_0)} \left( \frac{V^{(cell)}(s)}{V^{(cell)}(s_0)} \right)^{\frac{2}{3}}$ | $l_{AP}^{(cell)}(s_0) \sqrt[3]{\frac{V^{(cell)}(s)}{V^{(cell)}(s_0)}}$ | |
| Anisotropic growth only along $AP$ axis $l_{AP}^{(cell)}(s)$ | $A^{(bb)}(s_0)$ | $l_{AP}^{(cell)}(s)$ | |
| Convolution in the transverse plane | $\frac{V^{(cell)}(s_0)}{l_{AP}^{(cell)}(s_0) \gamma(s)}$ | $l_{AP}^{(cell)}(s_0)$ | |

#### Geometric modelling of cell neighbourhood shape changes

To compute how these geometric transformations affects the length of a group of cells, we define a measure of cell intercalation at each stage. To this aim, we assume that the length contribution of a group of *n* cells (where we assume that *n* is large) in the *AP* direction is given by *l*^(*n*)^ and the equation

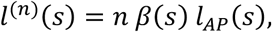

with the intercalation correction *β*(*s*). For *β* = 1, the cells are stacked on the AP axis, for small *β ≳* 0, many layers of cells are present on the DV and ML planes (see Fig. 2).

**Figure A2.**
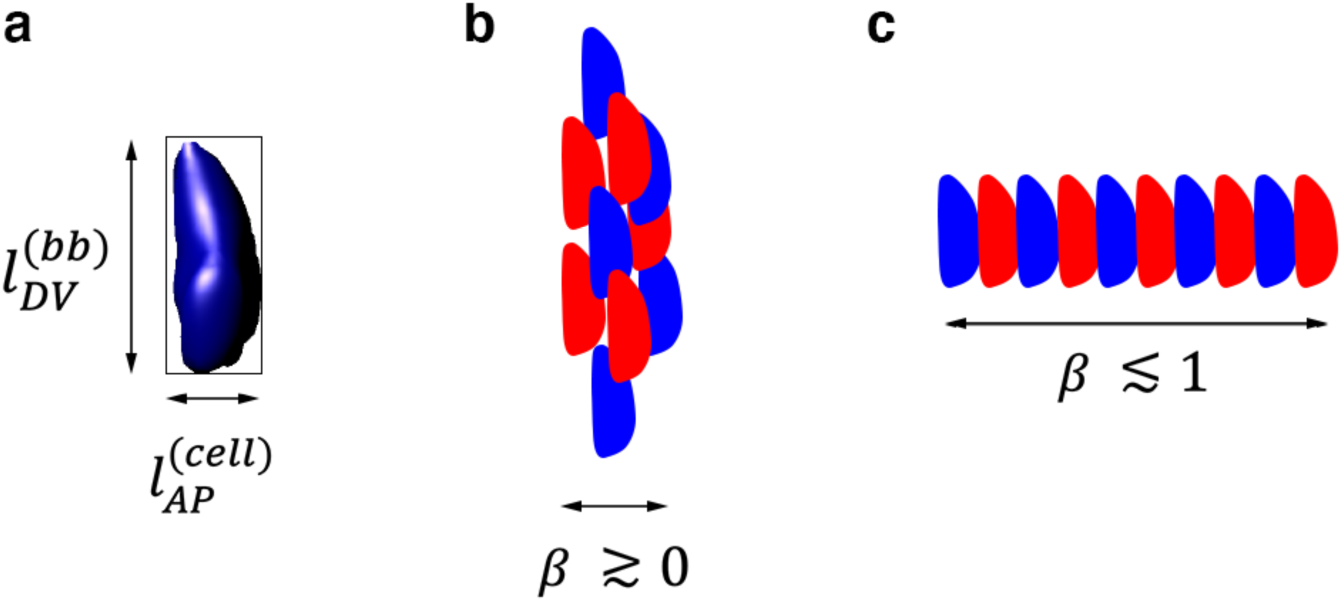
Calculation of intercalation correction *β* in a neighbourhood of 10 cells. (a) Sample cell from 8ss in lateral view showing the direction of AP length 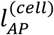, approximated by 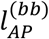. (b, c) Schematic of a neighbourhood of 10 cells in lateral view, showing two extremes of *β*, approaching 0 in (b) and approaching

We now start with a group of *n* cells, each of volume *V*^(*cell*)^(*s*), area *A*^(*cell*)^(*s*) and ratio *γ*(*s*), and an intercalation of *β*(*s*), and calculate neighbourhood length as

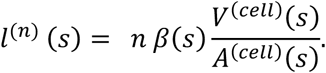

We can study the effect of each geometric transformation, with and without intercalation, on neighbourhood length *l*^(*n*)^ (*s*):

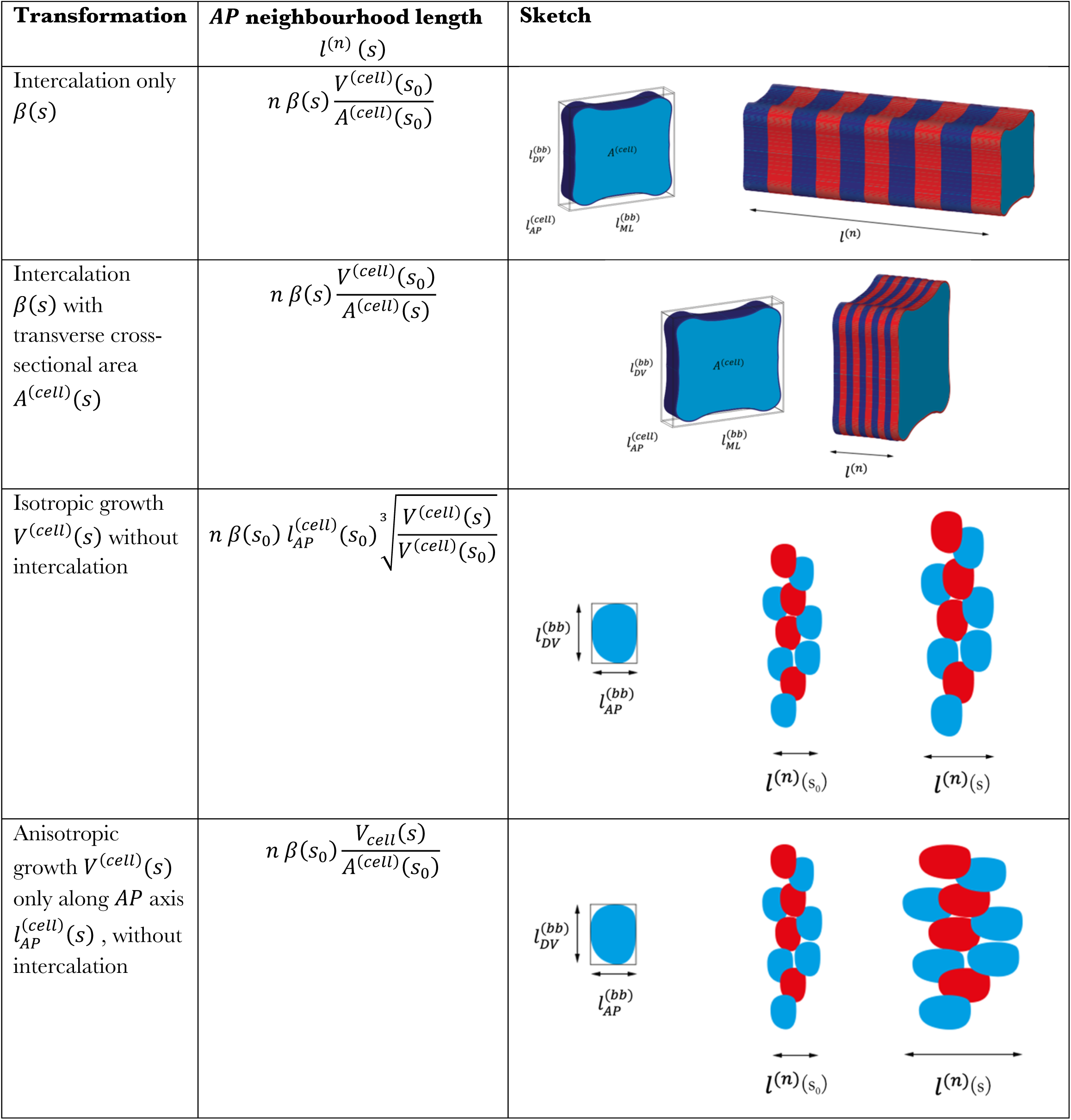

