## Supplement for "Single-cell morphometrics reveals ancestral principles of notochord development"

### Supplementary Information

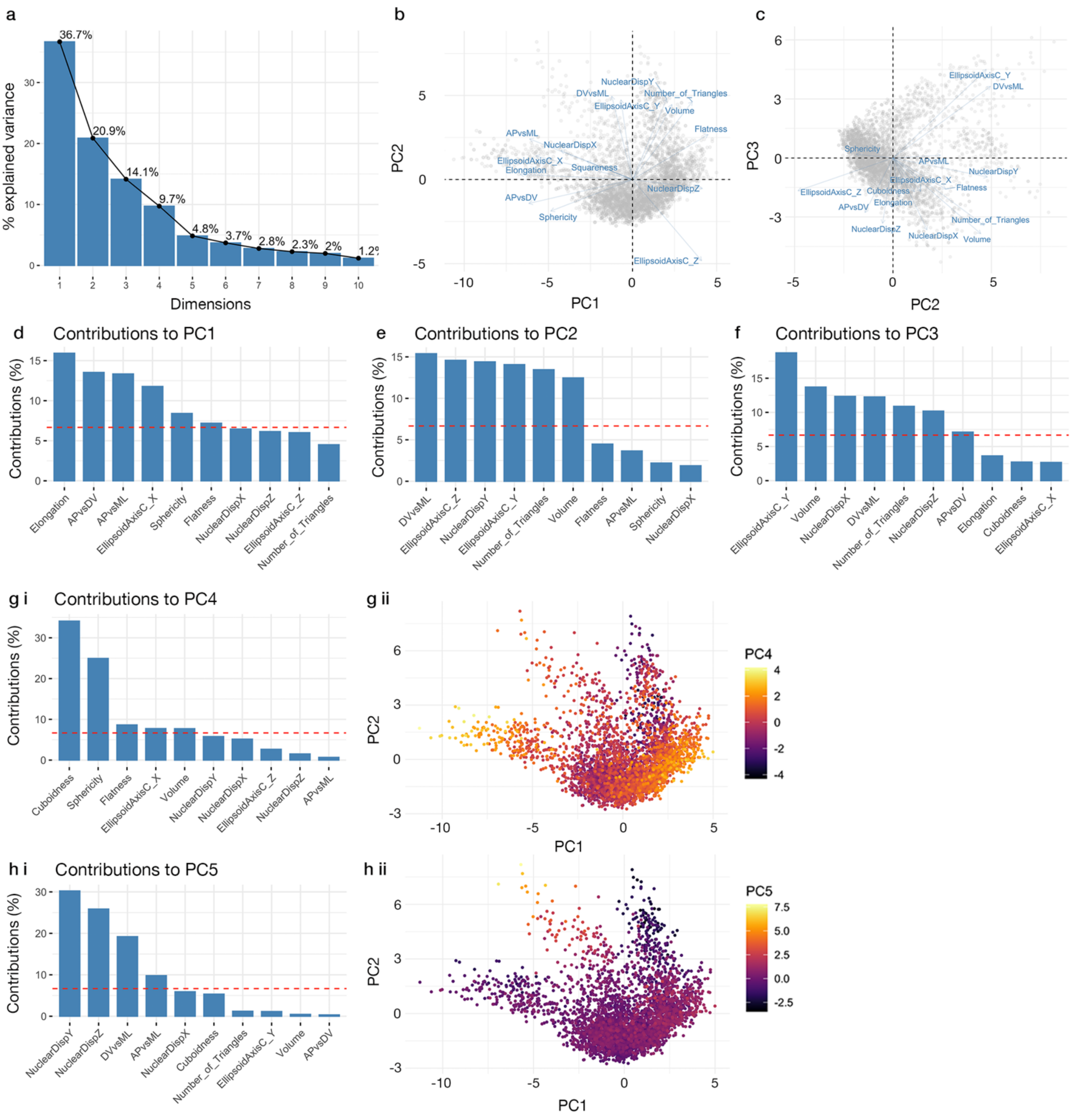


Figure S1. **Principal component analysis**. (a) Scree plot showing contribution of eigenvectors to total dataset variation. First 5 eigenvectors account for 86.2% total variation. (b, c) Compass plot showing direction of correlation between individual shape variables against PC1 and PC2 (b) and PC2 and PC3 (c). (d-h) Major correlates of PCs. f ii and g ii show all plots plotted against PC1 and PC2, colour-coded for PC4 (f ii) and PC5 (g ii).

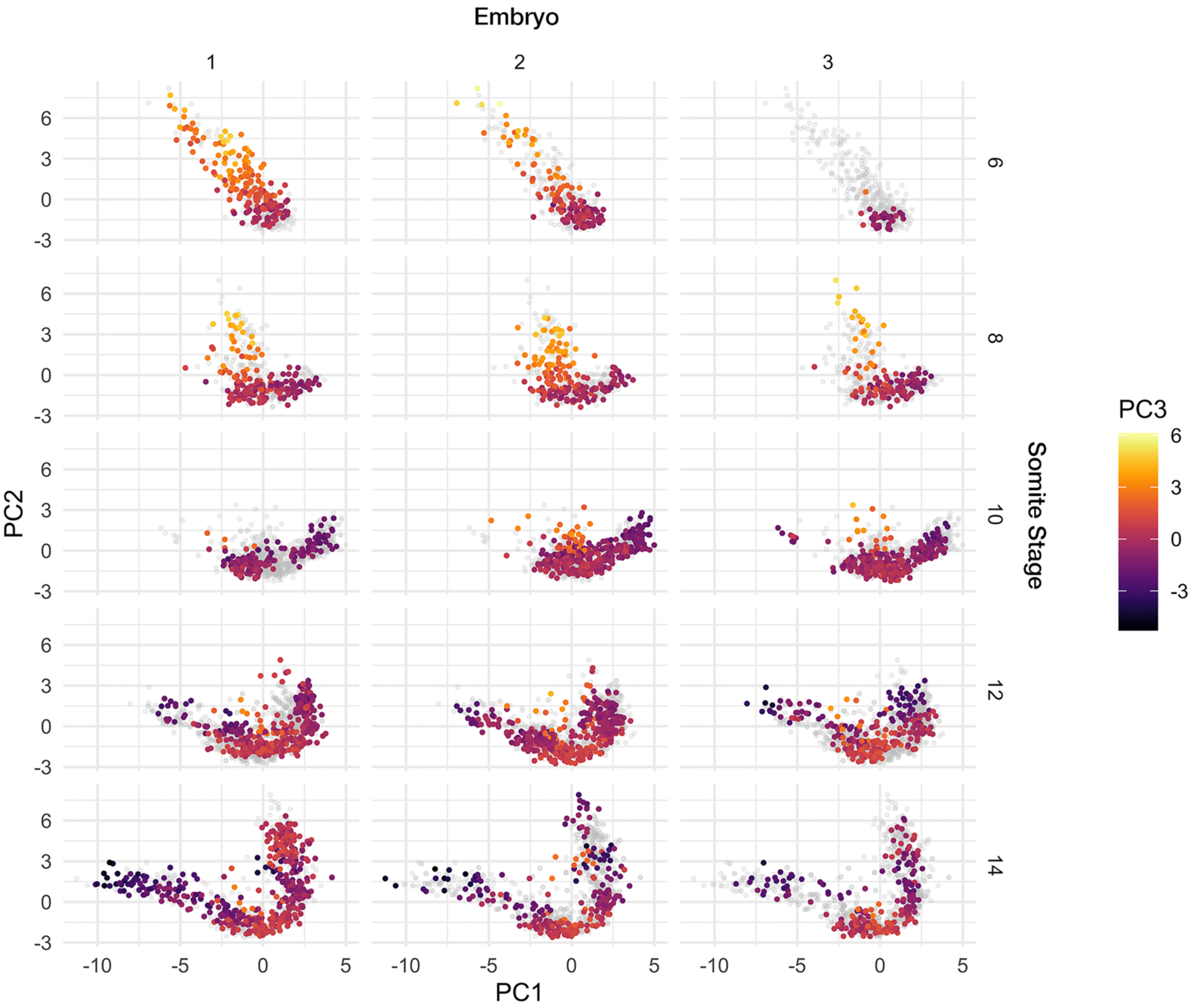


Figure S2. **Cell shape diversity is reproducible between different embryos.** Graphs show total morphospace filtered by somite stage and embryo. Grey points show all cells for the specified developmental stage. Colour-code illustrates distribution on PC3.

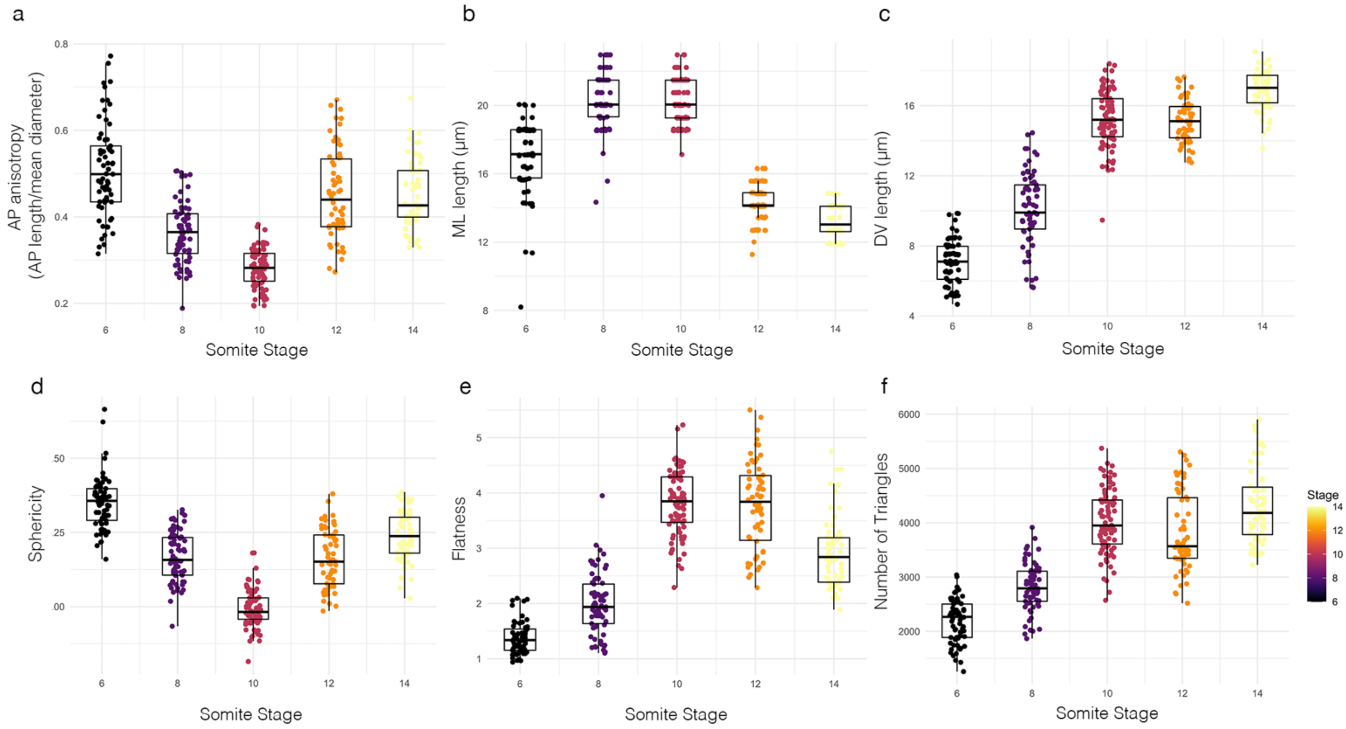


Figure S3. **Further geometric changes in cells from the 50% level of the notochord.** AP anisotropy (a), ML length (b), DV length (c), sphericity (d), flatness (e) and number of triangles (f).

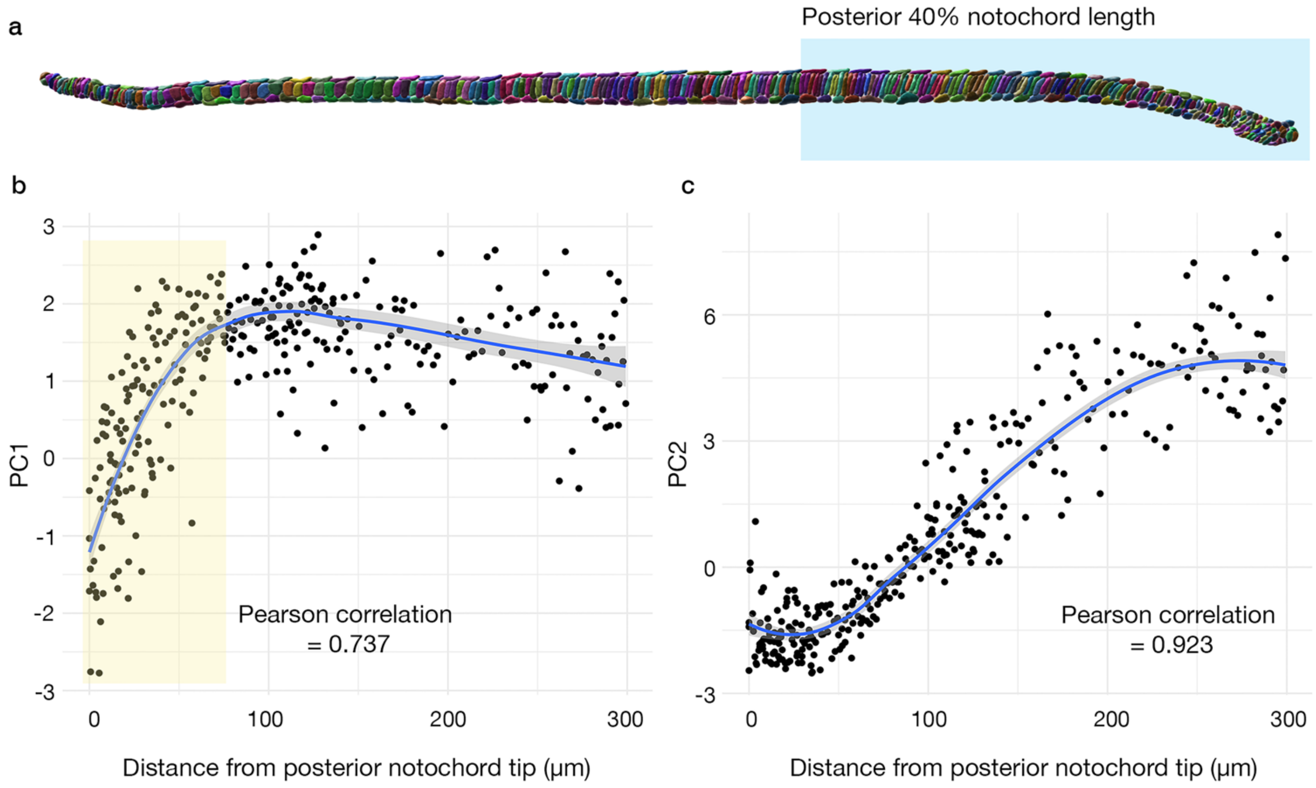


Figure S4. **Anteroposterior position correlates with developmental maturity in the posterior notochord**. (a) Notochord from the 14ss, with posterior 40% highlighted. (b) Graph shows AP position in the notochord plotted against PC1. Cells in the most posterior notochord (highlighted) exhibit a strong positive correlation between AP position and PC1, with a Pearson correlation coefficient of 0.737. (c) Graph shows AP position in the notochord plotted against PC2. Cells in this region exhibit a strong positive correlation between AP position and PC2, with a Pearson correlation coefficient of 0.923. Curves show a running mean and the standard error.

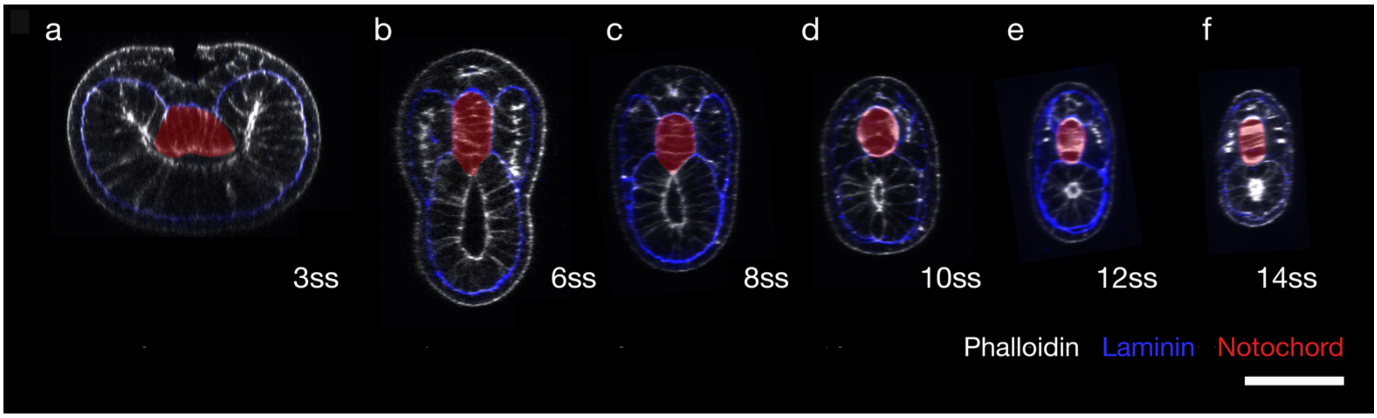


Figure S5. **Temporal changes in notochord morphology in transverse section.** Representative specimens from named stages across axis segmentation, in transverse section through the 50% level of the notochord. The notochord is highlighted in red. Specimens were stained for actin with phalloidin (white), to mark cell outlines, and immunostained for laminin (blue),

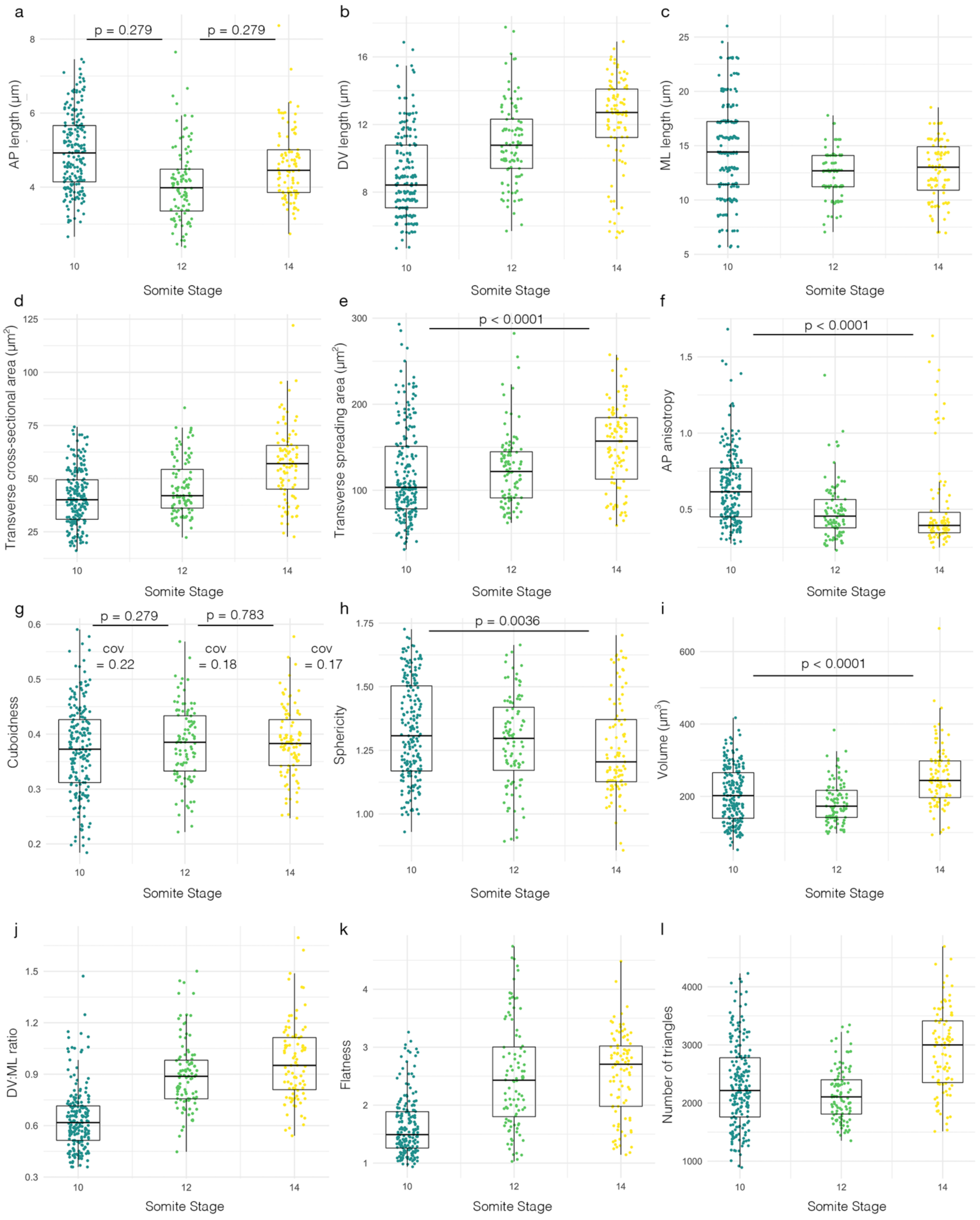


Figure S6. **Geometric changes in cells from the anterior tip of the notochord (anterior 15%).** (a-l) Graphs show geometric transitions between the 10ss and 14ss, when anterior cells are undergoing intercalation,; AP length (a), DV length (b), ML length (c), transverse cross-sectional area (d), transverse spreading area (e), AP anisotropy (f), cuboidness (g), sphericity (h), volume (i), DV:ML ratio (j), flatness (k), number of triangles (l). p values are the result of an unpaired Student's t-test between groups.

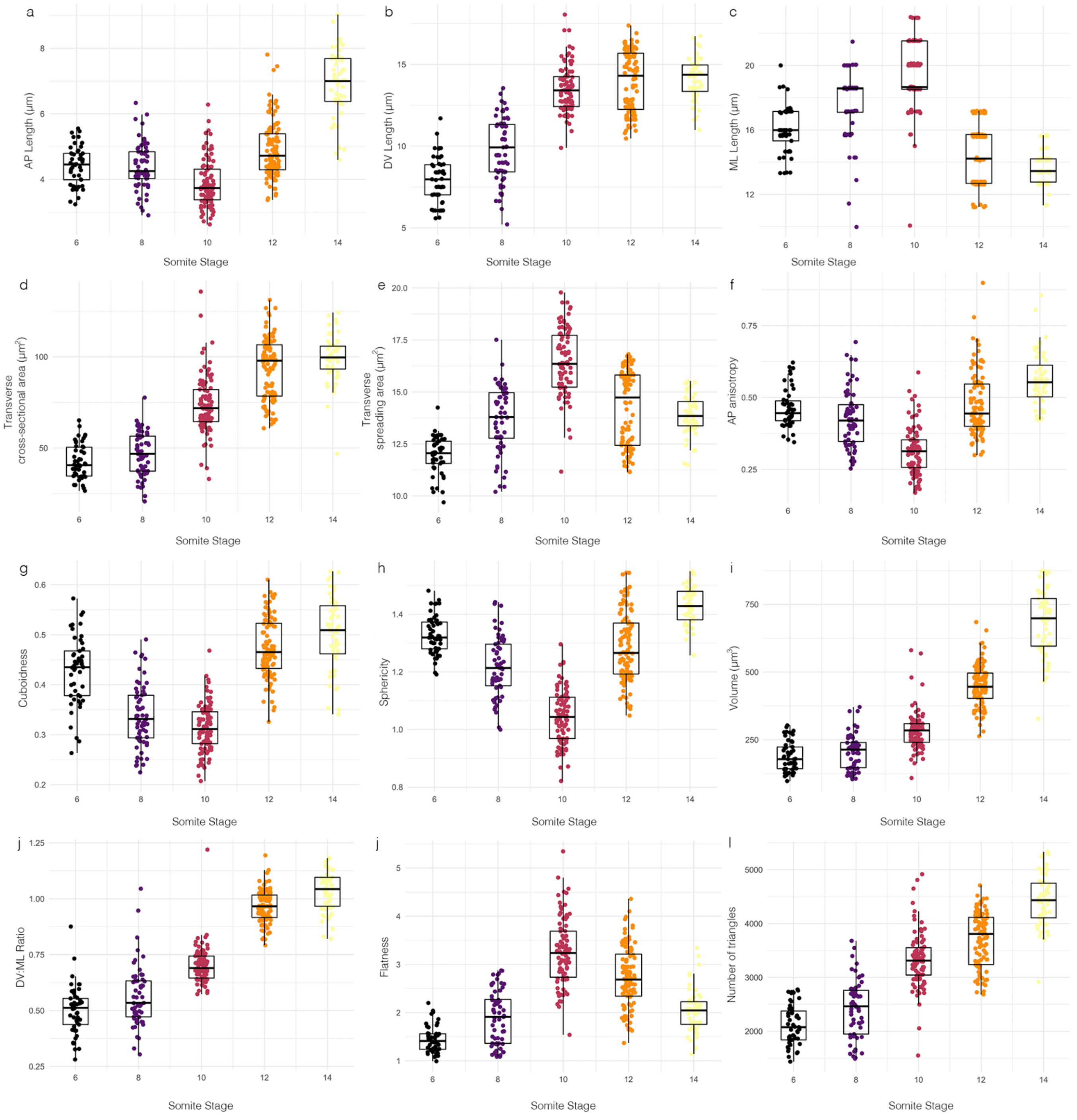


Figure S7. **Geometric changes in cells from the pharyngeal region of the notochord.** (a-l) Graphs show changes in AP length (a), DV length (b), ML length (c), transverse cross-sectional area (d), transverse spreading area (e), AP anisotropy (f), cuboidness (g), sphericity (h), volume (i), DV:ML ratio (j), flatness (k), number of triangles (l).

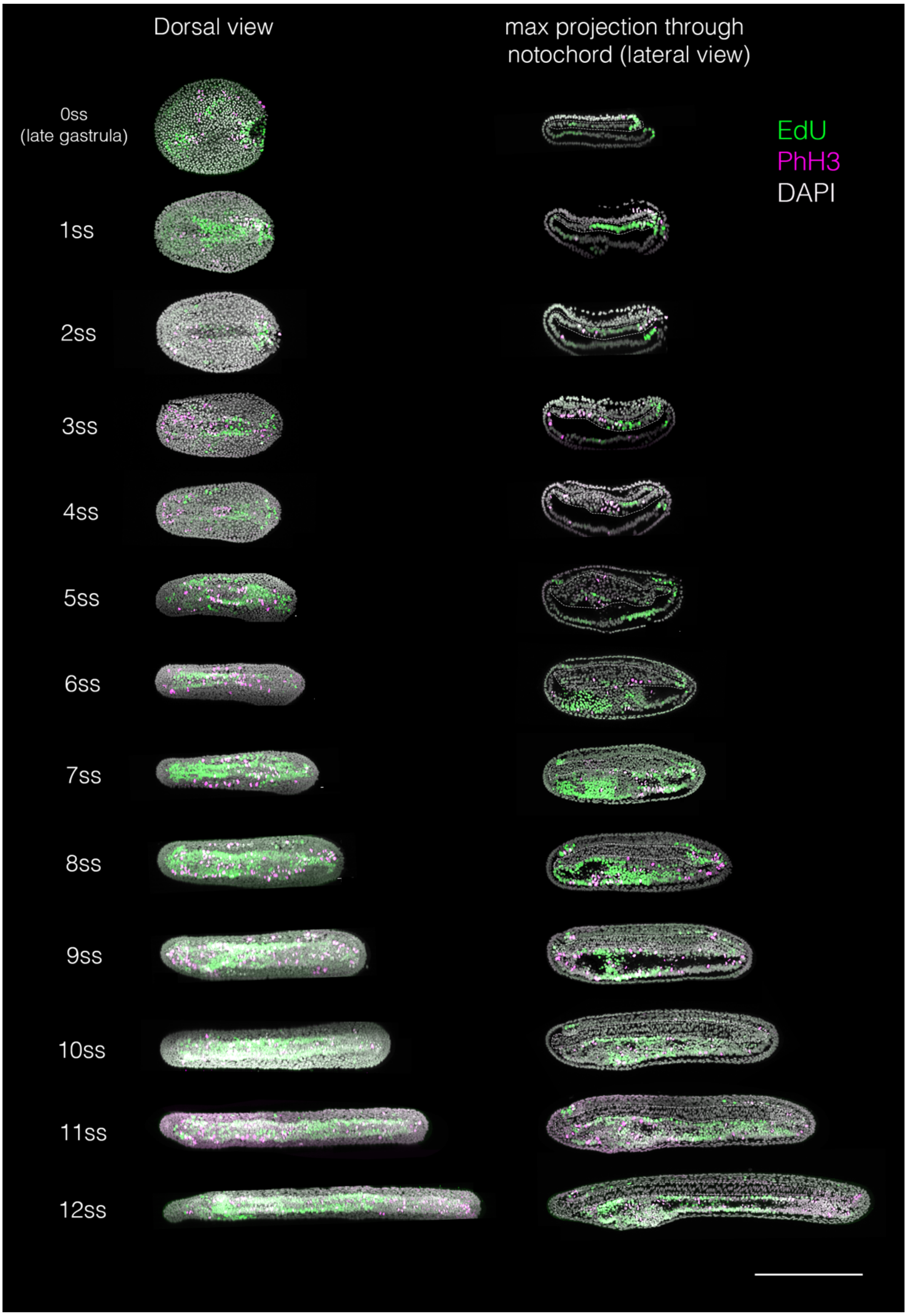


Figure S8. **Labelling for proliferative cells between gastrulation and 14ss.** Embryos at successive somite stages were incubated in 20μM EdU for 2 hours prior to fixation (green), and additionally immunostained for PhH3 after fixation (magenta). For each stage, embryos are in maximal projection in a dorsal view (left) and in midline sagittal section (right). Scale bar shows 200μm.
